# An *in vitro* platform for the enzymatic characterization of the rhomboid protease RHBDL4

**DOI:** 10.1101/2024.10.13.618094

**Authors:** Satarupa Bhaduri, Mac Kevin E. Braza, Stancho Stanchev, Marina Tauber, Raghad Al-Bawab, Lawrence J. Liu, Diego F. Trujillo, Kristina Solorio-Kirpichyan, Ambuj Srivastava, Javier Sanlley-Hernandez, Anthony J. O’Donoghue, Marius K. Lemberg, Rommie Amario, Kvido Strisovsky, Sonya E. Neal

## Abstract

Rhomboid proteases are ubiquitous intramembrane serine proteases that can cleave transmembrane substrates within lipid bilayers. They exhibit many and diverse functions, such as but not limited to, growth factor signaling, immune and inflammatory response, protein quality control, and parasitic invasion. Human rhomboid protease RHBDL4 has been demonstrated to play a critical role in removing misfolded proteins from the Endoplasmic Reticulum and is implicated in severe diseases such as various cancers and Alzheimer’s disease. Therefore, RHBDL4 is expected to constitute an important therapeutic target for such devastating diseases. Despite its critical role in many biological processes, the enzymatic properties of RHBDL4 remain largely unknown. To enable a comprehensive characterization of RHBDL4’s kinetics, catalytic parameters, substrate specificity, and binding modality we expressed and purified recombinant RHBDL4, and employed it in a Förster Resonance Energy Transfer-based cleavage assay. Until now, kinetic studies have been limited mostly to bacterial rhomboid proteases. Our *in vitro* platform offers a new method for studying RHBDL4’s enzymatic function and substrate preferences. Furthermore, we developed and tested potential inhibitors using our assay and successfully identified peptidyl α-ketoamide inhibitors of RHBDL4 that are highly effective against recombinant RHBDL4. We utilize ensemble docking and molecular dynamics (MD) simulations to explore the binding modality of substrate-derived peptides bound to RHBDL4.

Our analysis focused on key interactions and dynamic movements within RHBDL4’s active site that contributed to binding stability, offering valuable insights for optimizing the non-prime side of RHBDL4 ketoamide inhibitors. In summary, our study offers fundamental insights into RHBDL4’s catalytic activities and substrate preferences, laying the foundation for downstream applications such as drug inhibitor screenings and structure-function studies, which will enable the identification of lead drug compounds for RHBDL4.

## Introduction

Intramembrane proteases are evolutionarily widespread (found ubiquitously across all kingdoms of life, ranging from bacteria and archaea to eukaryotes) and play crucial roles in various biologically and medically relevant processes. Their unique localization within the lipid bilayer allows them to act as molecular scissors and cleave transmembrane-anchored substrates, which in turn has profound effects on downstream biological events (Wolfe 2009; Strisovsky 2013; Kühnle et al. 2019; Paschkowsky et al. 2019; Kandel and Neal 2020; Lemberg and Strisovsky 2021). Among the intramembrane proteases, rhomboid proteases are the best studied at the structural and molecular level (Düsterhöft et al. 2017). They are classified as serine proteases and function in growth factor signaling, parasitic invasion, inflammation, and immune response.

Rhomboid proteases belong to the rhomboid superfamily of proteins, which share a core six-transmembrane-helix bundle that is stabilized by the highly conserved WR motif and a Gx3G motif found in loop 1 (L1) and transmembrane helix 6 (TM6), respectively (Lemberg and Freeman 2007; Bergbold and Lemberg 2013; Tichá et al. 2018). Rhomboid-like proteins are generally classified as rhomboid intramembrane proteases and pseudoproteases. Specifically in mammals, there are five intramembrane proteases (rhomboid-like [RHBDL] 1–4, and presenilin-associated rhomboid-like [PARL]) and nine intramembrane pseudoproteases ([Derlin]-1–3, inactive rhomboid [iRhom] 1 and 2, UBA domain-containing [UBAC] 2, transmembrane protein [TMEM] 15, and rhomboid domain-containing [RHBDD] 2 and 3) (Adrain and Cavadas; Düsterhöft et al. 2017).

RHBDL4 has been well characterized at the molecular level and consists of six transmembrane helices, with a serine-histidine catalytic dyad between TM4 and TM6. It also contains the highly conserved rhomboid motifs WR and Gx3G, along with a p97/valosin-binding motif (VBM) and a ubiquitin interacting motif (UIM) at its C terminus, which suggests that the molecular function of RHBDL4 involves coordinated ubiquitin binding and p97/VCP recruitment (Fleig et al. 2012; Lim et al. 2016) (Fig. 1A). RHBDL4 is best characterized for its role in Endoplasmic Reticulum (ER) protein quality control, where it targets misfolded substrates for ER-associated degradation (ERAD) (Fleig et al. 2012; Kandel and Neal 2020). RHBDL4-triggered ERAD commonly involves the following processes: 1) ubiquitination of a misfolded substrate by an E3 ubiquitin ligase; 2) binding of misfolded substrate to RHBDL4 via its UIM; and 3) substrate cleavage of the misfolded substrate via the S144 protease site of RHBDL4. This is followed by recruitment of the AAA+ATPase p97/VCP, via the RHBDL4 VCP interacting motif, retrotranslocation of the cleaved protein fragments outside of the ER, and degradation by the cytosolic proteasome. The removal and degradation of misfolded substrates helps alleviate the proteotoxic stress within the ER(Fleig et al. 2012).

**Figure 1.**
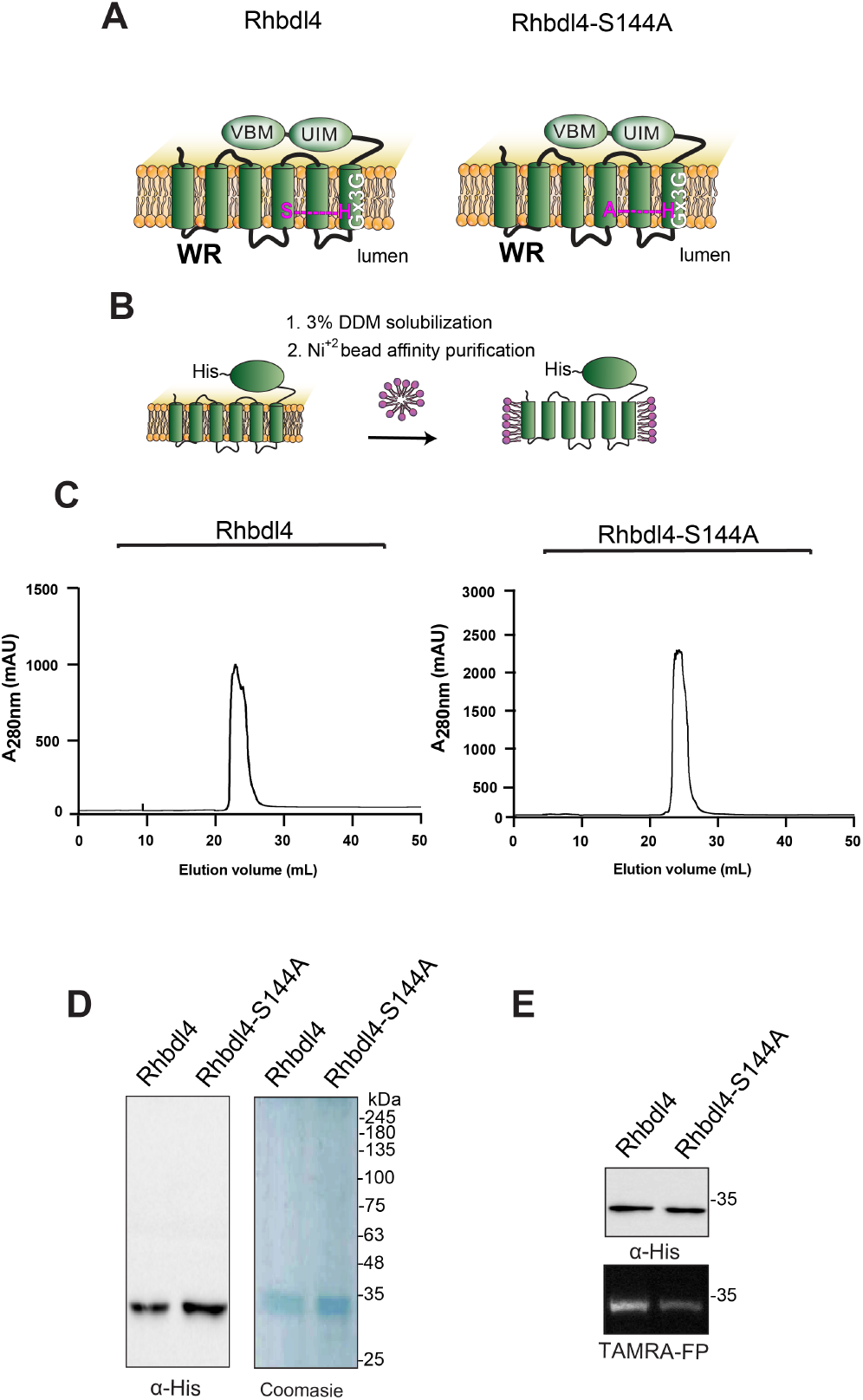
Purification of the rhomboid protease RHBDL4. **(A)** Schematic representation of the wild-type RHBDL4 and proteolytically inactive RHBDL4-S144A, highlighting their conserved rhomboid motifs (WR and Gx3G), VBM, and UIM domains. In the inactive RHBDL4, residue S144 is mutated to an alanine. (**B**) Schematic representation of RHBDL4 purification (His-tagged at its C terminus), solubilized in 3% DDM, and subjected to Ni^2+^ affinity purification. (**C**) Size-exclusion chromatography profile of wild-type RHBDL4 and inactive RHBDL4-S144A, being eluted as a monomeric protein. **(D)** Coomassie staining and western blotting with α-RHBDL4 antibody (Sigma) of concentrated purified wild-type RHBDL4 and inactive RHBDL4-S144A. **(E)** Purified wild-type RHBDL4 and inactive RHBDL4-S144A incubated with TAMRA-FP probe. The reactions were quenched with 1×Laemmli buffer and analyzed by SDS-PAGE on a 12% gel. Post-transfer membranes were visualized using a fluorescence Cy3 filter and subjected to western blotting using an α-RHBDL4 antibody.

RHBDL4 has been identified to play critical roles in both health and disease. On one side, RHBDL4 can cleave and degrade native proteins thereby regulating the activity of the (Oligosaccharyltransferase) OST complex (Knopf et al. 2020), or the trafficking factors for TMED7 (Knopf et al. 2024). On the other side, RHBDL4 specifically targets misfolded proteins such as a disease-associated mutant MPZ (L170R), which is linked to Charcot-Marie-Tooth disease, a condition marked by muscular weakness and wasting, as well as sensory loss (Fleig et al. 2012). Moreover, RHBDL4 can target and cleaves the APP ectodomain, leading to decreased Aβ peptide production, a component that is a hallmark of Alzheimer’s disease (Paschkowsky et al. 2016). Finally, RHBDL4 has been shown to regulate lipid biosynthesis through cleavage of SREBP1c (Han et al.). Additionally, the roles of RHBDL4 in disease have been studied in the context of cancer. RHBDL4 expression is upregulated in glioblastoma, colorectal, and liver cancer cell lines and is linked to tumor growth (Liu et al. 2013; Wei et al. 2014; Miao et al. 2017; Zhang et al. 2018). Clinical studies indicated that patients with low RHBDL4 expression in colorectal cancer tissue had better survival outcomes than those with higher RHBDL4 expression (Miao et al. 2017). Overall, a thorough examination of RHBDL4 function underscores its numerous and diverse roles in physiology and disease.

Although such functional diversity could make RHBDL4 a primary therapeutic target, its enzymatic properties, which are crucial for drug design, remain understudied. Here, we expressed and purified recombinant RHBDL4 and developed a Förster Resonance Energy Transfer (FRET)- based assay for *in vitro* cleavage to enable, for the first time, a complete characterization of its enzymatic properties and substrate specificity. First, we investigated the influence of pH and different salt concentrations on protease activity. RHBDL4 displayed excellent tolerance in a salt range concentration of 100–300 mM, with the highest activity being displayed at pH 8. Moreover, we found that the RHBDL4 conserved WR motif in the L1 region is required for its proteolytic activity. By contrast, mutating the conserved Gx3G motif did not abrogate RHBDL4’s proteolytic activity in our *in vitro* assay. Like GlpG and PARL, the RHBDL4 catalytic rate for proteolysis was slow with a cleavage rate of approximately 11/h. Additionally, our substrate specificity and docking studies revealed that, much like other rhomboid proteases characterized to date, RHBDL4 preferred small amino acids at the P1 position within its substrates (Strisovsky et al. 2009a; Zoll et al. 2014a; Tichá et al. 2017b; Bach et al. 2024). Finally, our *in vitro* platform set the stage for screening effective inhibitors against this critical rhomboid protease. As a proof-of-principle, peptidyl α-ketoamide inhibitors were derived from cleavage recognition sequence of known substrates along with another identified from the candidate screen in the study and were shown to inhibit RHBDL4 proteolytic activity. Overall, our *in vitro* platform will facilitate future drug discovery efforts for identifying RHBDL4 small-molecule modulators that can be further optimized to treat relevant diseases.

## Results

### Expression and purification of RHBDL4 and RHBDL4-S144A

RHBDL4 is an intramembrane protease, the catalytic residues of which are encoded by S144 and H195 (Fig. 1A and B). To generate a proteolytically inactive RHBDL4, the catalytic serine was mutated to an alanine (S144A) by site-directed mutagenesis. The entire RHBDL4 and RHBDL4- S144A cDNAs were subcloned into the pcDNA3.1 vector under the control of the CMV promoter, and C-terminally labeled with a 6×His tag. To evaluate the expression of RHBDL4 and RHBDL4-S144A, Freestyle^TM^ 293-F cells were transiently transfected with the corresponding plasmids, and their expression was evaluated on days 2, 3, 4, 5, and 6 after transfection by western blot analysis using an anti-His-tag antibody. We found that the level of expression for RHBDL4-S144A peaked on days 3-4 post-transfection, whereas RHBDL4 expression peaked on day 2 (Fig. S1A and & B). Thus, cells were harvested on day 2 for RHBDL4 and day 3 for RHBDL4-S144A for purification.

To produce large quantities of recombinant proteins for subsequent assays, Freestyle^TM^ 293-F cells expressing RHBDL4 and RHBDL4-S144A were harvested on day 2 or 3 post-transfection, and the membrane protease was solubilized in 3% DDM, purified using Ni^+2^ affinity chromatography, treated with TEV protease to release the affinity tag, and fractionated on size-exclusion chromatography (Fig. 1B). Following this, peaks with the highest absorbance values corresponded to fractions that were enriched with purified RHBDL4 or RHBDL4-S144A (Fig. 1C), as analyzed by SDS-PAGE, in which specific bands of purified RHBDL4 and RHBDL4-S144A were observed at approximately 34 kDa using western blotting and Coomassie staining (Fig. 1D). To confirm that the purified RHBDL4 is an active serine protease, active-site labeling was carried out by incubating 1 μg of purified RHBDL4 or inactive RHBDL4-S144A with the fluorescent serine hydrolase active-site labeler, TAMRA-FP. Notably, the fluorescent signal was reduced by ∼80% with inactive RHBDL4-S144A in comparison to active RHBDL4 (Fig. 1E). The cause of this residual activity by the catalytic mutant is unknown but similar behavior has been observed for the *E. coli* rhomboid GlpG (Sherratt et al. 2012). Collectively, our results confirmed the successful purification of recombinant rhomboid serine protease RHBDL4.

### Screening for peptide substrates cleaved by RHBDL4

We employed a FRET-based assay to determine the activity of RHBDL4. The RHBDL4 proteolytic activity was tested against a suite of fluorescent, internally quenched (IQ) substrates that have been previously used to test the activity of rhomboid protease PARL (Fig. S2A)(Lysyk et al. 2021). These substrates consist of 7–10-mer peptide sequences flanked by the fluorescent donor group 7-methoxycourmarin (mc) on the N terminus and quencher acceptor group dinitrophenol (dnp) on the C terminus (Fig. 2A) (Arutyunova et al. 2018). In general, the donor and acceptor are in proximity within the fluorogenic peptides and show low fluorescence due to internal quenching. Hydrolysis of the peptide bond between the donor and the acceptor leads to physical separation of the FRET pairs and increase in fluorescence, which would be proportional to RHBDL4 protease activity (Fig. 2A).

**Figure 2.**
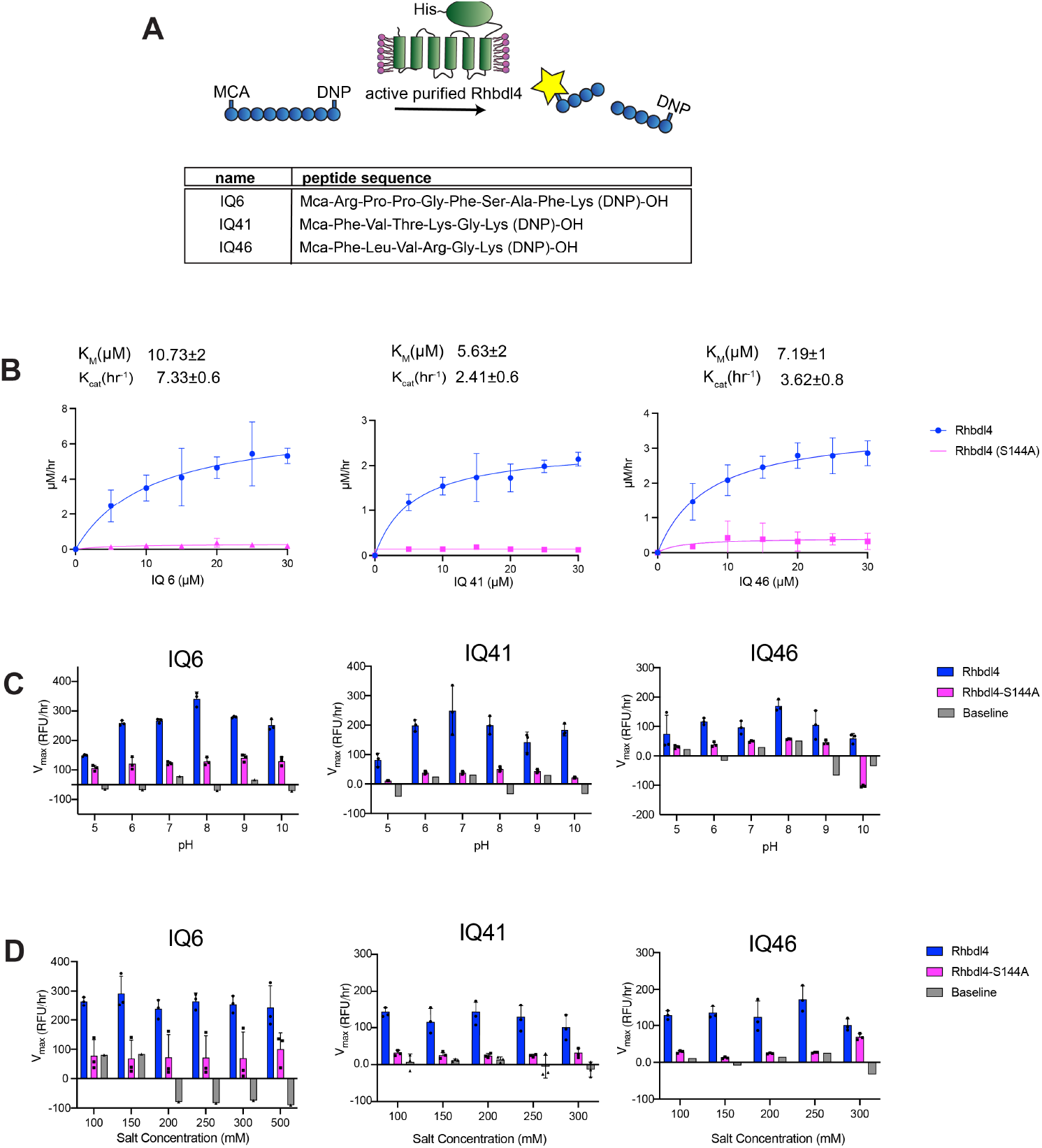
Kinetic characterization of recombinant human RHBDL4. (**A**) Schematic representation of the FRET-based cleavage assay using internally quenched (IQ) substrates. (**B**) Saturation curves of RHBDL4 using the fluorescent IQ substrates, IQ6, IQ41, and IQ46. (**C**) pH dependence of the RHBDL4 protease activity using substrates IQ6, IQ41, and IQ46. (**D**) Salt concentration dependence of the RHBDL4 protease activity using substrates IQ6, IQ41, and IQ46. All the assays were performed in triplicate, and data points are represented by the average ± standard deviation

### Kinetic analysis of RHBDL4

Incubation of freshly purified RHBDL4 with substrates IQ6 mca-RPPGFSAFK(dnp), IQ41 mca- FVTKGK(dnp), and IQ46 mca-FLVRGK(dnp) at pH 8 and 37°C (Fig. 2B) resulted in enzyme concentration-dependent increase in fluorescence (Fig. S2A), whereas a negligible increase in fluorescence was observed with purified inactive RHBDL4-S144A. The reaction setup in our FRET assay enabled the direct determination of Michaelis–Menten kinetic parameters K_m_ and k_cat_ for the cleavage of IQ6, IQ41, and IQ46 by RHBDL4. Our fluorescence reading data were converted to μM/h using the standard calibration curve for all three IQ substrates, and the correction factor for the inner filter effect was applied. We confirmed that purified RHBDL4 followed Michaelis–Menten kinetics with 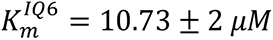, 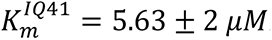 and 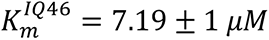 and 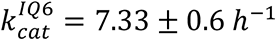, 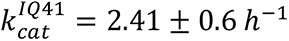, and 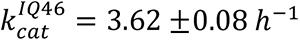 (Fig. 2B and S2B). Specifically, RHBDL4 exhibited three-fold and two-fold faster kinetics with IQ6 in comparison to IQ41 and IQ46, respectively. Thus, results from our FRET assay indicated that recombinant RHBDL4 was enzymatically active.

Our FRET-based assay is a suitable platform for optimizing conditions that enhance RHBDL4 protease activity. To this end, we selected the optimal pH for RHBDL4 protease activity and performed FRET-based assays with IQ6, IQ41, and IQ46 at a pH range of 5–10. RHBDL4 protease showed the highest activity with the IQ6 and IQ46 substrates at pH 8 and with the IQ41 substrate at pH 7 (Fig. 2C). The optimal salt concentration for the RHBDL4 protease activity was also tested in the range of 100–300 mM (Fig. 2D). We conclude that with the IQ6, IQ41, and IQ46 substrates, RHBDL4 had satisfactory activity in a salt concentration range of 100–300 mM.

### Conserved rhomboid motifs are required for RHBDL4 activity

Fig. 3A shows a schematic representation of RHBDL4 with its transmembrane (TM) domains highlighted in green, cytosolic protein domains highlighted in blue, and a list of specific motifs that were mutated. All mutants had robust expression on day 3 post-transfection of Freestyle^TM^ 293-F cells, when cells were harvested for purification. RHBDL4 contains two motifs that are well conserved among the rhomboid superfamily, the WR motif in loop 1 (L1) and the Gx3G motif in TM domain 6 (Fig. S3A and B) (Lemberg and Freeman 2007). Both motifs are required for RHBDL4-mediated ERAD function (Fleig et al. 2012). We predicted that both AR and Ax3G mutants would result in lower RHBDL4 activity. To that end, RHBDL4-AR and RHBDL4-Ax3G were expressed and purified using affinity Ni-NTA column followed by SEC (Fig. 3B and C). Purified recombinant RHBDL4, RHBDL4-AR, RHBDL4-Ax3G and control, inactive RHBDL4- S144A were employed in the FRET-based assay using IQ6 as the substrate. Wild-type and inactive RHBDL4-S144A showed the expected kinetics, whereas RHBDL4-AxR behaved similarly to the inactive RHBDL4-S144A and had a marked decrease (∼6-fold) in activity compared with the wild-type RHBDL4 (Fig. 3D). Notably, RHBDL4-Ax3G retained enzymatic cleavage activity at levels comparable to wild-type RHBDL4 (Fig. 3D). Altogether, the WR motif appears to play a critical role in RHBDL4 activity, whereas in the detergent-solubilized state the Gx3G motif function is not critical for core proteolytic activity and may be required for other functions of RHBDL4.

**Figure 3.**
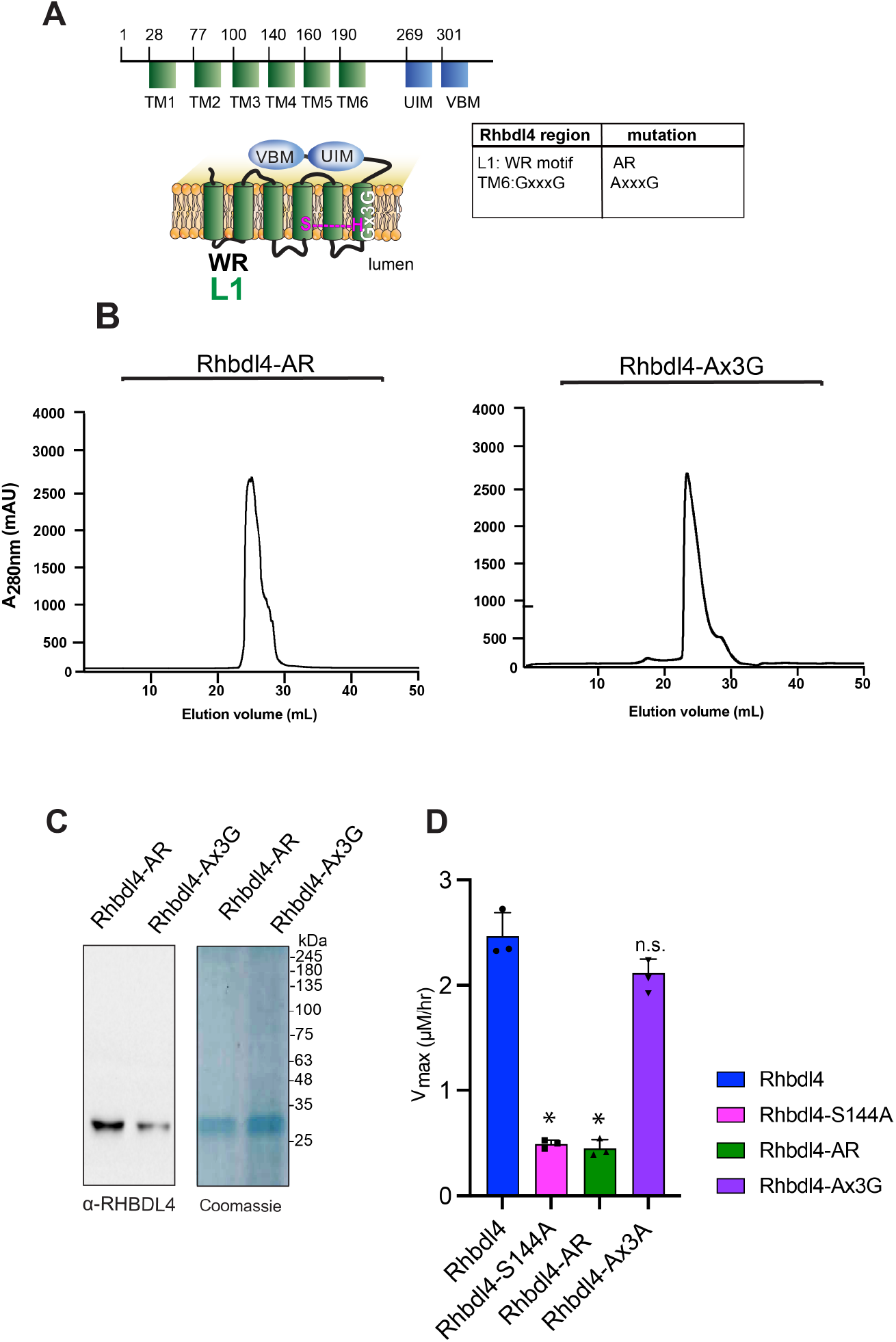
Dependence of conserved rhomboid motifs on RHBDL4 protease activity. **(A)** RHBDL4 representation highlighting TM1–6 (*in green*), the VBM and UIM binding motifs (*in blue*), the conserved rhomboid motifs (WR and Gx3G), and the serine-histidine catalytic dyad (*in pink*). The table indicates the RHBDL4 region, the conserved rhomboid motifs, and amino acid mutation. the RHBDL4 proteolytic activity. **(B)** Size-exclusion chromatography profile of RHBDL4-AR and RHBDL4-Ax3G, being eluted as a monomeric protein. **(C)** Coomassie staining and western blotting using an α-RHBDL4 antibody of concentrated purified RHBDL4-AR and RHBDL4-Ax3G. **(D)** Activity profiles of the wild-type RHBDL4 and mutant RHBDL4-S144A, RHBDL4-AR, and RHBDL4-Ax3G proteins, as measured by cleavage of the fluorescent substrate IQ6. Assay was performed in triplicate, and data points are represented by the average ± standard deviation.

### Substrate specificity of RHBDL4

Rhomboid proteases achieve substrate specificity based on the recognition motifs of their substrates and by reading poorly understood transmembrane domain dynamics, typically cleaving their substrates after a small amino acid (between the P1 and P1’ positions of substrates) (Strisovsky et al. 2009). We utilized mass spectrometry to identify RHBDL4 cleavage sites within the IQ6 substrate. We identified two N-terminal cleavage products, mca-RPPGFSA and mca- RPPGFS, after a 4-h incubation of RHDBL4 with IQ6 substrate mca-RPPGFSAFK(dnp) (Fig. S4A and B). This indicates that the protease preferentially cleaves the IQ6 substrate with Ala or Ser in the P1 position. Moreover, purified inactive RHBDL4-S144A has negligible activity towards IQ6, demonstrating that IQ substrates are specifically cleaved by RHBDL4 and not by other, potentially contaminating proteases. The bacterial rhomboid protease GlpG and mammalian rhomboid protease PARL have been previously shown to cleave the IQ4 substrate mca- RPKPYAvWMK(dnp) and PGAM-derived IQ15 substrate mca-AVFLSAVAG(dnp) respectively (Fig. S4C and D) (Arutyunova et al. 2018; Lysyk et al. 2021). We determined whether RHBDL4 can recognize and cleave both the fluorescent GlpG and PARL model substrates. We employed our *in vitro* cleavage assay but did not observe any RHBDL4-catalyzed activity towards IQ4 and IQ15 substrate (Fig. S4D). Taken together, our recombinant purified active RHBDL4 does not appear to bear specificity towards substrates that are recognized and cleaved by GlpG and PARL.

### In vitro inhibition studies of RHBDL4

To test whether the RHBDL4 rhomboid protease is sensitive to protease inhibitors, its activity was measured in the presence of general serine protease inhibitors such as N-*p*-tosyl-L-phenylalanine chloromethyl ketone (TPCK), 3,4-dichloroisocoumarin (DCI), and 4-(2-aminoethyl) benzenesulfonyl fluoride hydrochloride (ABESF). We found that TPCK and ABESF showed weak inhibition at all concentrations tested, whereas DCI inhibited the RHBDL4 proteolytic activity only at higher concentrations (100 mM) (Fig. 4A).

**Figure 4.**
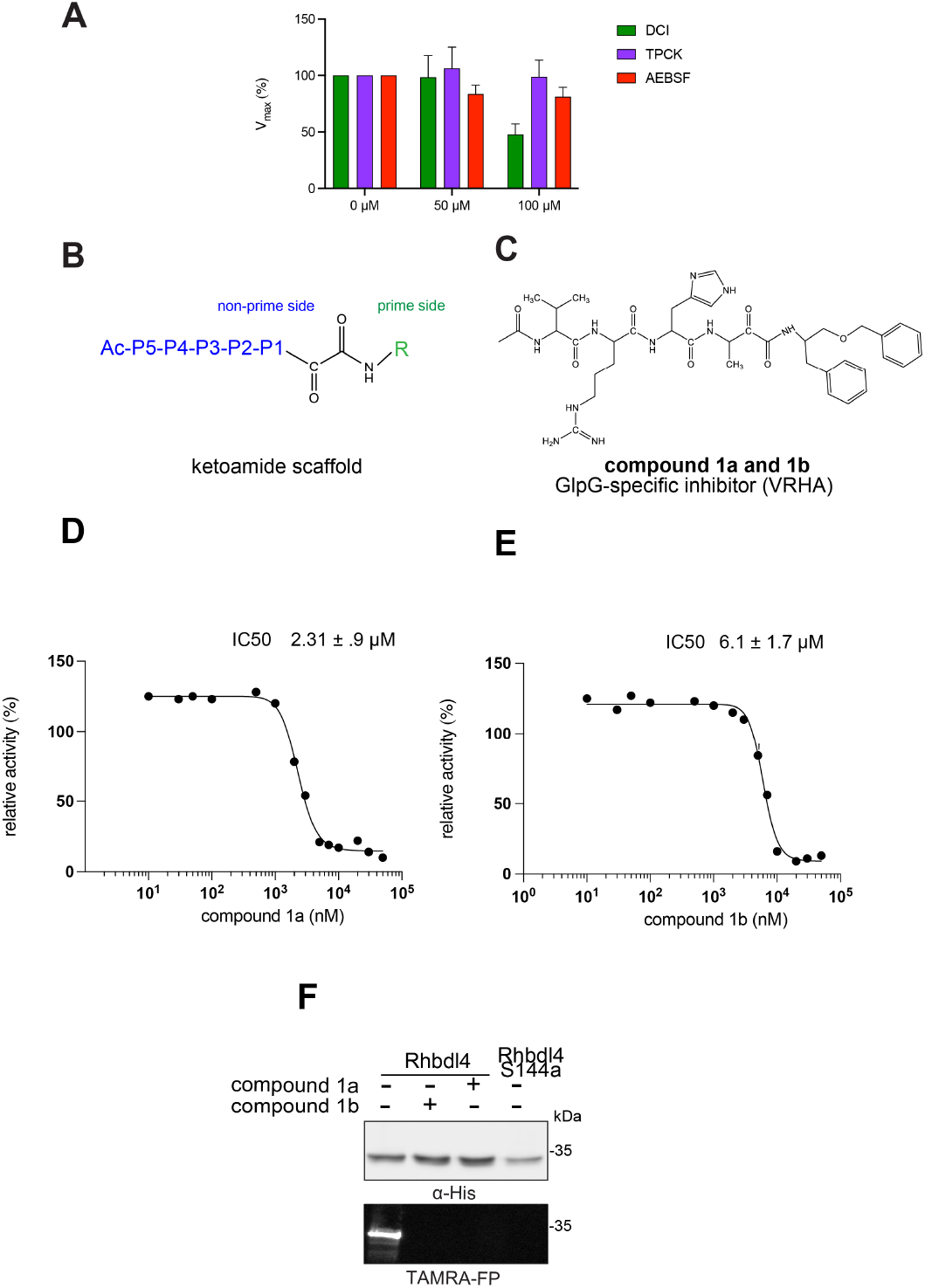
Inhibition of recombinant purified RHBDL4 proteolytic activity by N-substituted peptidyl α-ketoamides. (A) Proteolytic rates of the fluorescent substrate IQ6 by RHBDL4 in the presence of increasing concentrations of the general protease inhibitors, Dicyclocoumarin (DCI), tosyl-phenylalanine-chloromethyl ketone (TPCK) and 4-(2-aminoethyl) benzenesulfonyl fluoride (AEBSF). (B) Schematic representation of the peptidyl- α-ketoamide scaffold with peptide sequence P5-P1 at the non-prime side and hydrophobic substituent (R) at the prime side. (C) Schematic representation of the peptidyl- α-ketoamide inhibitor, compound 1a/1b. (D) Representative inhibition curve derived from measuring rates of RHBDL4 proteolysis of the fluorescent substrate IQ6 with increasing concentrations of compound 1a. (E) Same as (D), except inhibition curve was derived from using increasing concentrations of compound 1b. (F) Wild-type RHBDL4 and inactive RHBDL4-S144A were preincubated with TAMRA-FP probe followed by incubation with GlpG-specific peptidyl- α-ketoamide inhibitor 1a and 1b. Reactions were quenched with 1×Laemmli buffer and loaded on 12% SDS-PAGE gel. Post-transfer membranes were visualized using a fluorescence Cy3 filter and by western blotting using an α-RHBDL4 antibody. For (A), (D), and (E), all inhibitors were pre-incubated with RHBDL4 for 1hr and allowed to react with the IQ6 substrate for 2 h. Assays were performed in triplicate, and data points represent the average ± standard deviation.

Since the classical serine protease inhibitors did not exhibit robust efficacy, as expected (Strisovsky 2016; Tichá et al. 2017b), we next tested other potential inhibitors of RHBDL4 activity. Our *in vitro* assay provides a platform for leveraging medicinal chemistry to design, test, and identify inhibitors with high selectivity and potency against RHBDL4. Peptidyl α-ketoamides are promising candidates, as they mimic the substrate by interacting covalently and reversibly with the serine residue at the proteolytic site (Tichá et al. 2017). They are comprised of a preferred peptide sequence that can be derived from a substrate (typically spanning the P5/P4 to P1 positions) at the non-prime side and a hydrophobic ‘tail’ substituent (R) at the prime side (Fig. 4B). They have high potency and selectivity towards other rhomboid proteases including bacterial GlpG (Tichá et al. 2017), bacterial YqgP (Began et al. 2020), Plasmodium falciparum ROM4 (Gandhi et al. 2020), human PARL (Poláchová et al. 2023; Bach et al. 2024, and RHBDL2 (Bach et al. 2024) rhomboid proteases, and their mechanism was shown to be slow binding and non-competitive (Tichá et al. 2017; Poláchová et al. 2023).

Based on conceptual similarity, we have initially focused on a model a peptidyl α- ketoamide inhibitor, compound **1** (Fig. 4C and Table 1), which was originally designed for the *E.coli* GlpG based on its preferred peptide sequence (Tichá et al. 2017b) at the cleavage site, and which incorporates a branched substituent at the ketoamide nitrogen (Bach et al. 2024). We have generated isomers of compound 1 (designated **1a** and **1b**) that most likely differ in their configuration of the Cα atom on the His in the P2 position. Notably, using an in vitro activity assay with fluorogenic substrates, compounds **1a** and **1b** inhibited RHBDL4 with IC50 values of 2.31 ± 0.9 *μM* and 6.1 ± 1.7 *μM*, respectively (Fig. 4D & E). As anticipated, both peptidyl α-ketoamides were able to compete with a general active site probe for serine hydrolases, TAMRA-FP, which resulted in a significant reduction of fluorescent signal, a low intensity signal that is comparable to the inactive RHBDL4-S144A control (Fig. 4F). Given that the IQ6 substrate mca-RPPGFSAFK(dnp), identified through substrate candidate screening, exhibited the highest cleavage rate for RHBDL4 proteolysis, we proceeded to develop an IQ6-derived peptidyl α-ketoamide, compound **2** (Fig. S5A and Table 1), based on the P5-P1 peptide cleavage recognition sequence PGFSA. The in vitro assay showed that compound **2** also inhibited RHBDL4 activity with IC50 value of 3.4 ± 1.2 *μM*(Fig. S5B). Overall, our in vitro FRET-based assay provides a suitable platform for designing potential inhibitors for RHBDL4. Here, we demonstrated that GlpG-derived and IQ6-derived peptidyl α-ketoamides inhibited recombinant RHBDL4’s activity.

**Table 1.**
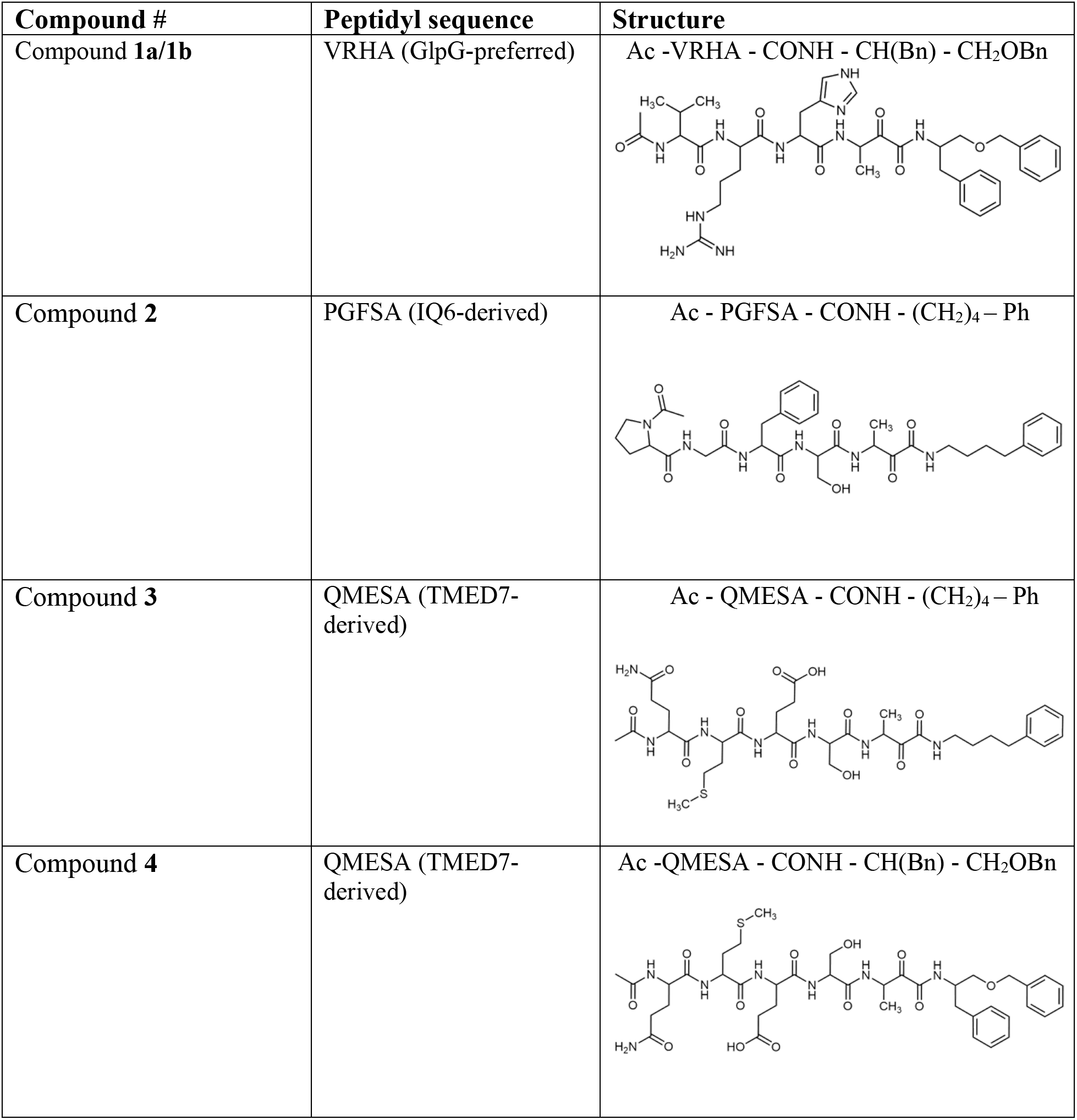
Peptidyl α-ketoamides used in this study.

| Compound # | Peptidyl sequence | Structure |
| --- | --- | --- |
| Compound **1a/1b** | VRHA (GlpG-preferred) | Ac -VRHA - CONH - CH(Bn) - CH<sub>2</sub>OBn |
| Compound **2** | PGFSA (IQ6-derived) | Ac - PGFSA - CONH - (CH<sub>2</sub>)<sub>4</sub> - Ph |
| Compound **3** | QMESA (TMED7-derived) | Ac - QMESA - CONH - (CH<sub>2</sub>)<sub>4</sub> - Ph |
| Compound **4** | QMESA (TMED7-derived) | Ac -QMESA - CONH - CH(Bn) - CH<sub>2</sub>OBn |

Recently, RHBDL4 has been shown to target and cleave TMED7, a secretory factor required for trafficking toll-like receptor 4 to the cell surface, which is a major mediator of innate immunity (Knopf et al. 2024). We leveraged our in vitro FRET-based activity assay to develop and test the activity of TMED7-derived peptidyl α-ketoamides. Branched substituents at the prime side of the compound (Fig. 4B, *highlighted in green*) have been shown to increase selectivity and potency towards some rhomboid proteases (Bach et al. 2024). Furthermore, previous mapping of the major cleavage site in TMED7 producing N-terminal cleavage fragment (P5-P1) ending with the sequence QMESA (Knopf et al. 2024). We used this substrate-derived sequence to produce a peptidyl α-ketoamide inhibitor equipped with a phenylbutyl (compound **3**) or branched phenylbutyl tails (compound **4**) tail added at the prime side (Fig. 5A and Table 1).

**Figure 5.**
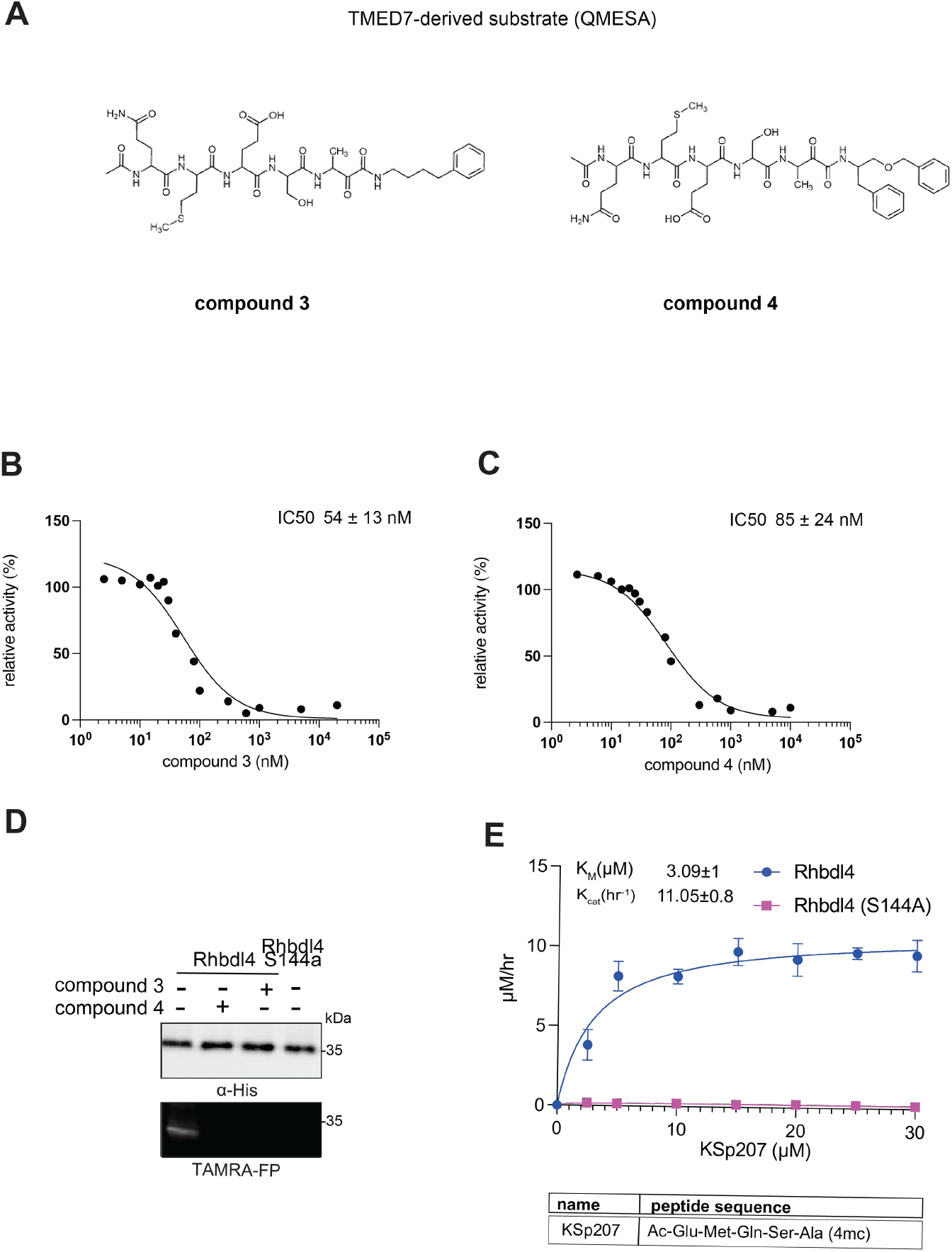
*In vitro* inhibition of RHBDL4 activity with TMED7-derived peptidyl α- ketoamides. **(A)** Schematic representation of the peptidyl- α-ketoamide inhibitor, compound **3** and **4. (B)** Representative inhibition curve derived from measuring rates of RHBDL4 proteolysis of the fluorescent substrate IQ6 with increasing concentrations of compound **3**. **(C)** Same as (B), except inhibition curve was derived from using increasing concentrations of compound **4**. **(D)** Wild-type RHBDL4 and inactive RHBDL4-S144A were preincubated with TAMRA-FP probe followed by incubation with peptidyl-α-ketoamide inhibitor **3** and **4**. Reactions were quenched with 1×Laemmli buffer and loaded on 12% SDS-PAGE gel. Post-transfer membranes were visualized using a fluorescence Cy3 filter and by western blotting using an α-RHBDL4 antibody. **(E)** Saturation curve of RHBDL4 using fluorescent substrate, KSp207. For (B) and (C), all inhibitors were pre-incubated with RHBDL4 for 1hr and allowed to react with the IQ6 substrate for 2 h. All assays were performed in triplicate, and data points represent the average ± standard deviation.

Notably, in the *in vitro* assay, both compounds inhibited RHBDL4 activity with similar IC50 values in the nanomolar range, 54 ± 13 *nM* and 85 ± 24 *nM* respectively (Fig. 5B & C). In this case, adding a branched substituent on the prime side did not influence the potency of these inhibitors, which has proven to be the case also for human RHBDL2 (Bach et al. 2024).

Moreover, both compounds were able to compete with the TAMRA-FP probe for RHBDL4’s catalytic site, causing a reduction in overall fluorescent signal, which resulted in low intensity signal that was comparable to the inactive RHBDL4-S144A control (Fig. 5D).

Thus far, TMED7-derived α-ketoamides have demonstrated the greatest *in vitro* potency in comparison to the other compounds tested, indicating that the TMED7 peptide sequence binds more effectively to RHBDL4’s active site. We were curious to see if a TMED7-derived fluorescent substrate would exhibit high cleavage activity by RHBDL4. We generated fluorescent substrate Ac-QMESA-4mc (KSp207), which contains the peptide sequence derived from the cleavage site in TMED7 (Knopf et al. 2024). Interestingly, KSp207 displayed higher enzymatic cleavage efficiency than IQ6, with 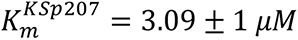 and 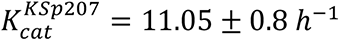 Fig. S5C), indicating that RHBDL4 prefers the TMED7 cleavage site sequence, which is consistent with the inhibition data.

### Inhibition studies of RHBDL4 in ER-enriched microsomes

So far, our ketoamide inhibitors exhibited a wide range of IC50 values in vitro (Table 1). We aimed to investigate our RHBDL4 inhibitors in a more physiological context. Since we did not observe any inhibition in the cell-based cleavage RHBDL4 assay, even at high concentrations of all compounds (100 μM) (data not shown), we used a TAMRA-FP competition label assay in ER-enriched microsomal fractions that increases accessibility to compounds but maintains RHBDL4’s native assembly with various ERAD factors (Bock et al. 2022). To achieve this, we prepared ER-enriched microsomes from HEK293T cells transfected with RHBDL4-GFP, then pre-incubated microsomes with 100 μM of each compound, before TAMRA-FP was added to the microsomal fractions. RHBDL4-GFP was immunoprecipitated and visualized for TAMRA-FP fluorescence signal and western blotting. Consistent with the *in vitro* assay using detergent solubilized and purified RHBDL4, compounds **3** & **4** reduced, and compound **1** almost completely inhibited TAMRA-FP labeling (Fig. 6). We surmise that the peptidyl α-ketoamides may have low membrane penetrability, making them weak inhibitors in microsome and cell-based assays. Further refinement is needed to design peptidyl α-ketoamides that effectively penetrate lipids and inhibit RHBDL4 activity in its native context. This is in principle possible, as other peptidyl ketoamides do inhibit GlpG (Bach et al. 2024)and PARL (Poláchová et al. 2023; Bach et al. 2024) in cells in their native membranes.Hence, our *in vitro* platform offers an efficient basis for the development of a new generation of RHBDL4 inhibitors that will be active in cells.

**Figure 6.**
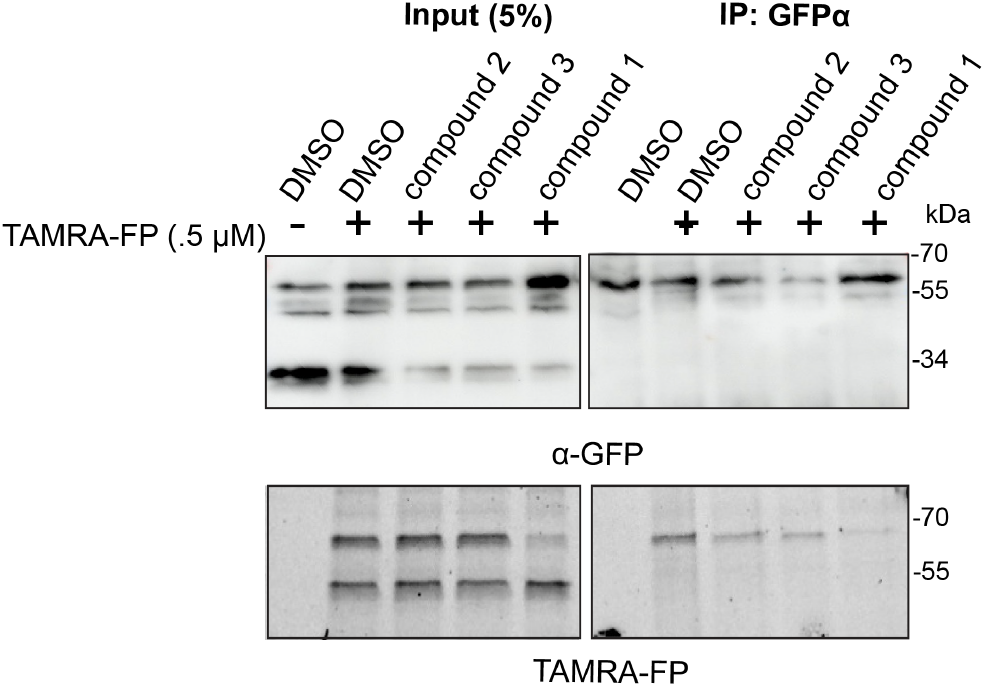
RHBDL4 inhibition studies in ER-enriched microsomes. ER-enriched microsomes were preincubated with 100 μM of compounds or DMSO vehicle control followed by addition of 500 nM of TAMRA-FP serine hydrolase probe. RHBDL4-GFP was immunoprecipitated with a α-GFP antibody and loaded on SDS-PAGE. TAMRA-FP labeling was analyzed via Cy3 filter followed by western blotting using α-GFP antibody.

### Substrate docking and molecular dynamics simulations

Given that rhomboid proteases exhibit broad yet distinct preferences for sequences from positions P5 to P1, we examined the dynamic interactions between substrate-derived peptides and the active site of RHBDL4 at the atomic level. Accordingly, molecular docking and explicitly-solvated all atom molecular dynamics (MD) simulations were used to probe the structural effects of substrate-derived peptide binding to RHBDL4 (Fig. 7A & B). An AlphaFold 2 structure of the human RHBDL4 was generated (AF-Q8TEB9-F1-v4)(Jumper et al. 2021). The AF2 model was protonated at pH 7.0, placed into a lipid bilayer with ER-like composition, explicitly solvated with a water and ion buffer, and subjected to 4 replicas of 500 ns production dynamics (total 2 μs dynamics). The ER lipid bilayer was composed of 47% POPC, 20% POPE, 11% POPI, 7% POPS, and 15% cholesterol, based on the average membrane composition previously reported (van Meer et al. 2008; Casares et al. 2019). After dynamics, we clustered the resulting structures and obtained the most populated conformation of the APO form RHBDL4 from all-atom MD simulations, we used the most populated conformer that exhibited an accessible active site, which was used for ensemble molecular docking (of putative substrates) and additional substrate-bound MD (Fig. 7 and S6) (Amaro et al. 2018). Following alignment, substrate-derived peptides and RHBDL4 exhibited low root mean square deviation (Fig S7). We calculated the binding energies of PGFSA and QMESA substrates via AutoDock Vina, and both peptides showed comparable binding affinity to the RHBDL4 active site (Table S2). Based on ensemble docking, we identified RHBDL4’s residues that interacted with the peptide substrates via hydrophobic and hydrogen bonding interactions (Fig. 7C & S8, Table 2). In comparison to PGFSA, the QMESA peptide exhibited additional hydrophobic interactions with RHBDL4 TM5 residues I165 and A182. Moreover, the QMESA substrate P5 glutamine residue (Q) formed two extra hydrogen bonds with RHBDL4’s TM5 residue F186, further stabilizing the interaction (Fig. S8 & Table 2). These interactions are not observed in the PGFSA. Because the addition of P5 and P6 residues to the α-ketoamide inhibitors have been shown to enhance their potency in some cases (Cho et al. 2016), we investigated whether eliminating the P5 residue from QMESA (resulting in MESA) would destabilize the initial binding to RHBDL4’s active site. In comparison to QMESA, the MESA peptide did not exhibit hydrophobic interactions and hydrogen bonding with RHBDL4’s I165/A182 and F186 residues respectively. To assess whether the peptide substrates remained stably bound throughout the simulation, we monitored the distance evolution between the peptide substrates—using the P2 serine as a reference point for all three peptides—and the RHBDL4 catalytic residues (H195 and S144). Indeed, we observed unbinding of MESA from the active site after 450 ns (Fig. S9) suggesting that the initial binding pose of MESA is not stable. In contrast, both PGFSA and QMESA remained close to their initial binding pose, suggesting a stable and sustained interaction within the active site of RHBDL4 (Fig. 7D, S10, & S11).

**Figure 7.**
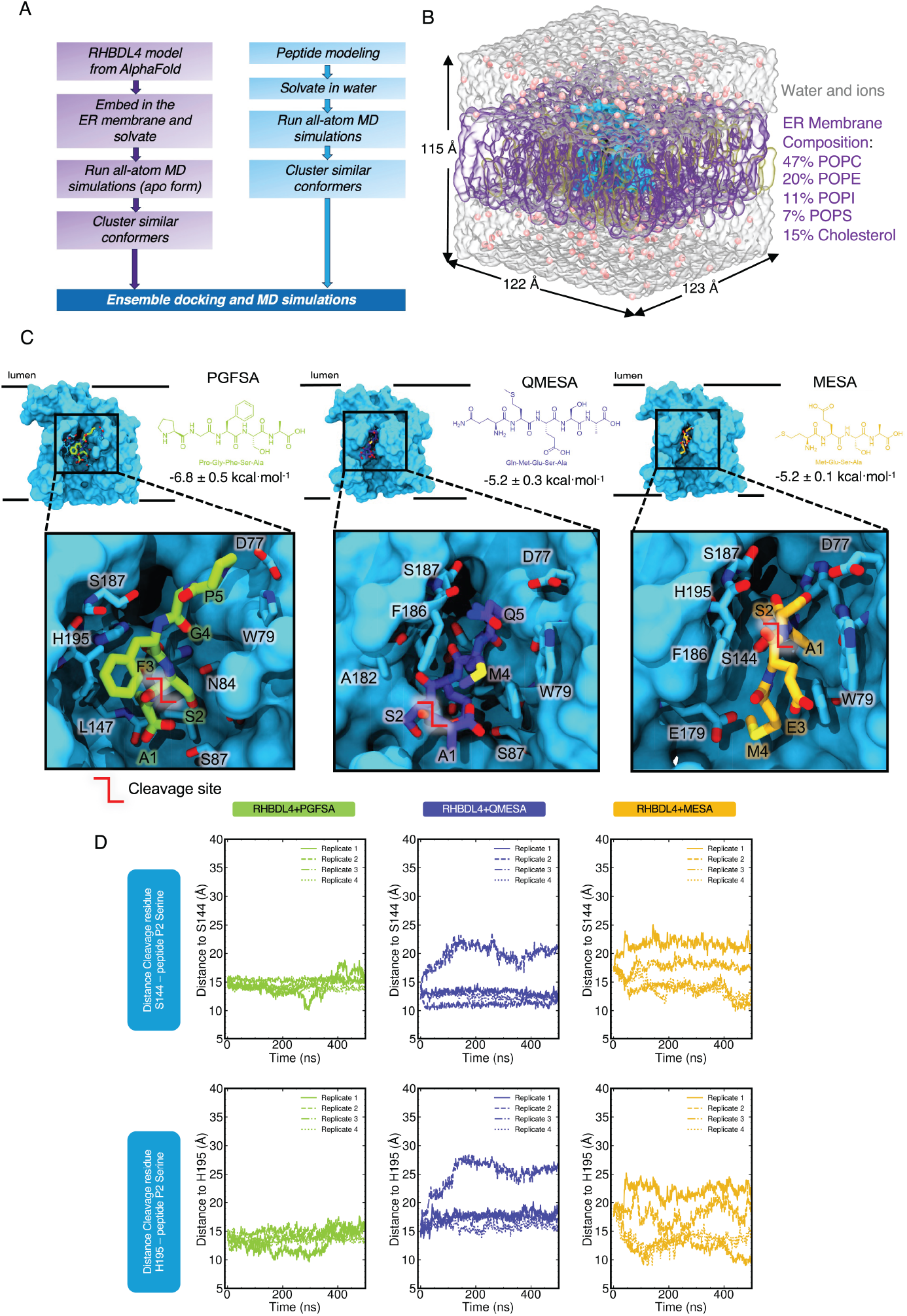
RHBDL4 structure modeling and peptide ensemble docking. RHBDL4 structure modeling and peptide ensemble docking. (A) Workflow of modeling RHBDL4 structural dynamics and docking of peptide substrate. Human RHBDL4 model was obtained from AlphaFold 2 and was equilibrated with all-atom MD simulations. Ensemble docking was done in AutoDock Vina with peptide (n=4) and RHBDL4 clustered structures from MD simulations. (B) Membrane composition and box size dimension. (C) Peptide structures, binding site, and calculated ensemble average binding energy. RHBDL4 open structure was used for the ensemble docking with the crevice adjacent to the catalytic dyad S144 and H195 exposed. (D) P2 residue and active site distance evolution.

**Table 2.** RHBDL4 Close interacting residues with the substrate. Refer to Supplemental Figure 8 for the 2D rendering of interaction.

| Peptide | Close interacting amino acid residues |  |
| --- | --- | --- |
|  | Hydrophobic interactions | Hydrogen bonds |
| PGFSA | D77 (TM2)<br>W79 (TM2)<br>H80 (TM2)<br>F83 (TM2)<br>I183 (TM5)<br>F186 (TM5)<br>S187 (loop 5) | E179 (TM5) |
| QMESA | D77 (TM2)<br>W79 (TM2)<br>H80 (TM2)<br>F83 (TM2)<br>I165 (TM5)<br>E179 (TM5)<br>A182 (TM5)<br>I183 (TM5)<br>S187 (loop 5) | F186 (TM5) |
| MESA | D77 (TM2)<br>W79 (TM2)<br>H80 (TM2)<br>F83 (TM2)<br>N84 (TM2)<br>E179 (TM5)<br>I183 (TM5)<br>F186 (TM5)<br>S187 (loop 5) | none |

We pinpointed regions of RHBDL4 that promoted peptide substrate accessibility. Notably, there were increased localized dynamic motions within RHBDL4’s active site following substrate binding. Specifically, the binding of PGFSA, QMESA, and MESA led to increased RMSF values (more than twice as mobile compared to the APO form of RHBDL4) in TM5, loop 5 (L5) and TM6 (Fig. 8A). The increased flexibility in L5 and TM6 affected the active site by increasing the distance between catalytic dyad residues, S144 and H195 (Fig. 8B). Moreover, binding of all substrate-derived peptides increased L5-TM2 distance characterized by residue pairs H80-S187 and H80-T190. Notably, amongst all the peptides, PGFSA induced the greatest increase by over 6 Å (Fig. 8C-F). This indicates that, even after substrate binding, the active site continues to enhance its accessibility, ensuring the substrate is precisely positioned for efficient proteolysis. Furthermore, this dynamic behavior aligns with previous observations of lateral gate openings in related rhomboid proteases, such as GlpG and PARL (Wu et al. 2006; Baker et al. 2007; Zoll et al. 2014b; Shokhen and Albeck 2017; Bohg et al. 2023).

**Figure 8.**
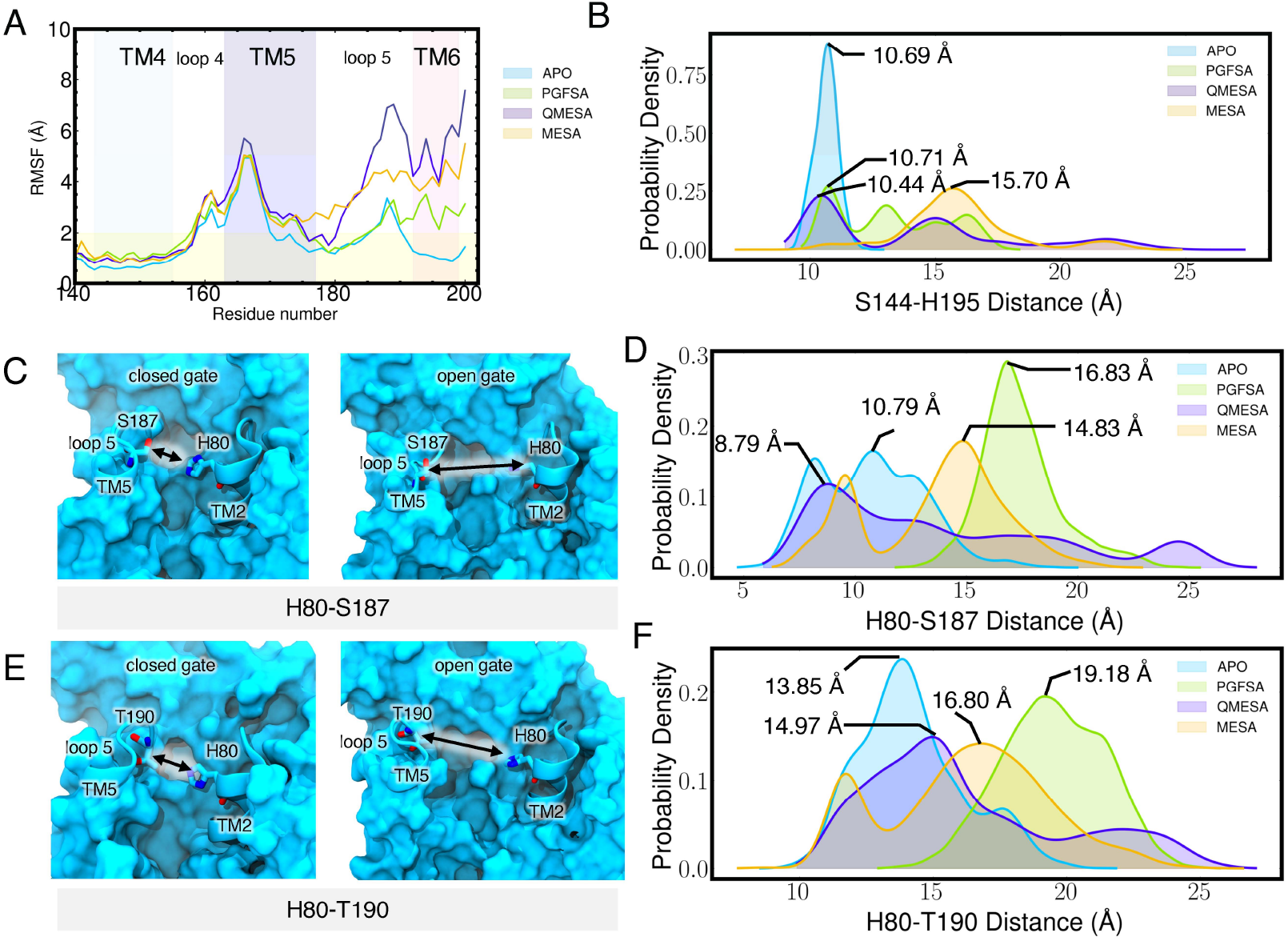
Structural effect of peptide binding to the RHBDL4. (A) RMSF calculated per residue for each simulated system. Calculated RMSF per residue were averaged over all replicates. (B) Active site catalytic dyad, S144 and H195 distance distribution. Distance were measured between the center of mass of residue S144 and H195. (C) H80-S187 distance during the closed and open gate.

## Technical limitations of this study

The MD analyses conducted in this study exclude the involvement of the ketoamide moiety of inhibitors, which is responsible for establishing covalent binding to the catalytic residue of RHBDL4 and the prime side substituent that contributes to the binding energy. Previous modifications to this region were shown to enhance potency and selectivity of ketoamide inhibitors against rhomboid proteases, including GlpG, PARL, and RHBDL4 (Bach et al. 2024) These limitations should be considered when interpreting the MD results. Nonetheless, the MD findings reveal important initial binding interactions between the peptide sequence and RHBDL4’s active site, which does strongly influence ligand binding to rhomboid (Bach et al. 2024) and serve as valuable insights for inhibitor design.

## Discussion

The development of *in vitro* cleavage assays has facilitated the study of rhomboid protease enzymatic activities outside of their complex cellular environment (Lemberg et al. 2005; Strisovsky et al. 2009; Dickey et al. 2013; Arutyunova et al. 2018, 2021; (Pierrat et al. 2011; Tichá et al. 2017a; Gandhi et al. 2020a; Bach et al. 2024). These platforms have enabled the identification and development of rhomboid protease inhibitors (Tichá et al. 2017a; Poláchová et al. 2023; Bach et al. 2024), marking the first step toward future therapeutic interventions.

RHBDL4 has garnered particular attention due to its contribution to biomedically important pathways (Bhaduri et al. 2022). Despite being a promising target for drug inhibitor development, its enzymatic function remains poorly understood. To address this gap, we expressed, purified, and developed an *in vitro* cleavage assay for RHBDL4. Our FRET-based *in vitro* assay not only allows for the exploration of RHBDL4’s kinetic activity and its substrate preferences but also facilitates the development and testing of potential RHBDL4 inhibitors, which hold significant therapeutic potential. Our results show that, like other studied rhomboid proteases, RHBDL4 has a slow cleavage rate. Utilizing our *in vitro* assay and substrate docking studies, we demonstrated that RHBDL4 exhibits a clear preference for substrates with an Ala residue in the P1 position, a preference similar to that of bacterial rhomboid protease GlpG (Tichá et al. 2017) and human rhomboid protease PARL (Poláchová et al. 2023; Bach et al. 2024). We also identify peptidyl α- ketoamide inhibitors with high potency against RHBDL4’s *in vitro* activity, which can serve as lead compounds for future drug design. Finally, ensemble docking and MD simulations of substrate-derived peptides bound to membrane-embedded RHBDL4 revealed that the P5 position residue plays a critical role in binding stability, offering practical insights for inhibitor design.

The RHBDL4 protease activity has primarily been characterized through cell-based assays. It possesses two critical motifs that are widely conserved in the rhomboid superfamily: the WR motif in its L1 loop region and the Gx3G motif in its TM domain 6 region (Fleig et al., 2012). Both motifs have been shown to be essential for the RHBDL4 ERAD function in cells (Fleig et al., 2012; Bock et al. Cell Reports, 2022). However, it remains unclear whether these motifs are necessary for RHBDL4’s core proteolytic activity. Mutation in the WR motif resulted in a significant decrease in RHBDL4’s enzymatic activity, consistent with earlier studies on rhomboid proteases such as YqgP, which found that a similar L1 WA mutant within the protease exhibited reduced cleavage activity (Lemberg et. Al, EMBO J., al. 2005). In contrast, disrupting the Gx3G rhomboid motif did not eliminate RHBDL4’s proteolytic cleavage activity in the detergent solubilized state. The role that the Gx3G motif plays in RHBDL4-mediated ERAD remains an important question.

Rhomboid proteases generally exhibit substrate specificity, which is imparted both by structural instability in the transmembrane segment of the substrate, and by the amino acid sequence around the site of cleavage, which influences both the cleavage site and rate (Strisovsky et al. 2009b, 2009a; Zoll et al. 2014a; Lysyk et al. 2021). Rhomboid proteases exhibit loose but recognizable preferences for the P5 to P1 sequences where the residue at the P1 position, immediately preceding the scission bond, is one of the strongest and common determinants (Strisovsky et al. 2009a; Zoll et al. 2014a). Several studies have highlighted the distinct substrate preferences of rhomboid proteases. For instance, bacterial rhomboid proteases such as AarA, GlpG, YqgP, along with human rhomboid protease PARL, strongly prefer substrates with small amino acid residues with non-branched side chains at the P1 position (Strisovsky et al. 2009; Zoll et al. 2014a; Saita et al. 2017; Arutyunova et al. 2018). Our mass spectrometry analysis reveals that RHBDL4 protease shares a similar substrate preference, favoring small amino acid residues like Ser and Ala at the P1 position within the IQ6 substrate. This aligns with recent studies, reporting RHBDL4’s preference for small residues at the P1 position in substrates MHC202 and TMED7 (Bock et al. 2022; Knopf et al. 2024). Despite these widespread observations, some rhomboid proteases can tolerate even large hydrophobic amino acid residues at the P1 position (Strisovsky et al. 2009b; Lysyk et al. 2021). Therefore, future studies focused on identifying the full range of amino acid residues that are preferred at the P5 to P1 positions in substates/inhibitors, including non-natural ones, and their possible interactions and synergies will be invaluable to the field.

In the past decade, there has been a notable surge in the development of rhomboid protease inhibitors. Initially, several inhibitors were identified, such as isocoumarins, β-lactams, β-lactones, and chloromethyl ketone derivatives (Tichá et al. 2018). However, these inhibitors bach had limitations; they lacked selectivity and were not effective as cell-based tools. A breakthrough came when peptidyl α-ketoamides were identified as particularly effective and selective inhibitors of rhomboid proteases. Importantly, these inhibitors targeted the active site in a substrate-like, reversible, and non-competitive manner (Began et al. 2020; Bach et al. 2024). Since then, peptidyl α-ketoamides have aided the functional characterization of several prokaryotic and eukaryotic rhomboid proteases (Began et al. 2020; Gandhi et al. 2020b; Poláchová et al. 2023; Bach et al. 2024). Our *in vitro* FRET-based assay has provided a platform for measuring the inhibition of rhomboid protease RHBDL4. We tested a suite of peptidyl α- ketoamides, including GlpG-derived, IQ6-derived, and TMED7-derived peptidyl α-ketoamides. We found that recombinant RHBDL4 has a clear preference for TMED7-derived inhibitor where residues from P5-P1 (QMESA) exhibited a strong inhibitory activity with a IC50 value in the nanomolar range. This is an exciting finding, given the role of RHBDL4 in a plethora of medically relevant pathways. We also attempted to test the efficacy of these inhibitors in isolated ER-enriched microsomes and in cells. However, we observe weak inhibition of RHBDL4 in ER-enriched microsomes and no inhibition in cells, suggesting these compounds may have low membrane penetrability or are inaccessible to RHBDL4’s active site in its native context. Nonetheless, our FRET-based *in vitro* assay is a powerful platform for further refining, testing, and identifying inhibitors that will be effective in inhibiting RHBDL4 *in vivo*.

MD simulations allowed us to observe the dynamic interactions between peptide substrates and membrane-embedded RHBDL4. In our *in vitro* FRET-based assay, the TMED7- derived peptidyl α-ketoamide inhibitor (QMESA) was more potent than the IQ6-derived inhibitor (PGFSA). However, our initial docking and simulation studies do not fully explain why QMESA exhibited greater potency. One possibility is that, beyond the time scale of the simulation, additional residues may be involved in further stabilizing QMESA within the active site compared to PGFSA. How the P5 residue within α-ketoamide inhibitors contributes to potency has been widely debated (Cho et al. 2016; Tichá et al. 2017). Our observations revealed that QMESA remains stably bound within the active site, whereas the variant MESA lacking the P5 residue disassociates from the active site. This difference is reinforced by the presence of additional hydrophobic interactions and hydrogen bonds with the P5 glutamine residue (Q), which effectively anchor the peptide within the active site of RHBDL4. Furthermore, we unexpectantly observed localized dynamic movements within RHBDL4’s active site even after substrate binding. These movements create space for substrates to orient and position themselves correctly near RHBDL4’s cleavage residue. The observed flexibility likely accounts for RHBDL4 active site’s ability to accommodate structurally diverse substrates. A previous study suggested that the lateral gate opening of the active site of bacterial rhomboid member GlpG, was the rate-limiting step for protease enzyme activity (Dickey et al. 2013). However, we observed that the APO RHBDL4 active site remained accessible and open over a timescale of several hundred nanoseconds to accommodate a substrate. This raises the intriguing possibility that, the rate-limiting step may not be the lateral gate opening but rather the correct positioning of the substrate within the active site. This opens an important avenue for future research to determine whether this is a unique feature of RHBDL4 or extends to other distant members of the rhomboid protease family (Tichá et al. 2017a, 2017b).

In summary, we expressed, purified, and evaluated the protease activity of recombinant rhomboid protease RHBDL4. Similar to other rhomboid proteases, the activity of RHBDL4 is pH dependent (Lemberg et al. 2005a; Tichá et al. 2017a), albeit weakly. RHBDL4 kinetic analysis revealed a slow catalytic rate, with the IQ6 and KSp207 substrate and cleavage occurring at a rate of approximately 7/h and 11/h respectively. According to our FRET substrate-based assay, RHBDL4 has a similar substrate preference profile to most rhomboid proteases characterized to date, which prefer small amino acid residues at the P1 cleavage position.

Furthermore, peptidyl α-ketoamides were found to exhibit inhibitory activity against RHBDL4. Overall, our findings not only further our understanding of RHBDL4 protease function, but also provide a basis for designing specific substrates and inhibitors.

## Author Contributions

S.B., M.K.B., L.J.L., D.T., K.S., A.J.O, R.O., M.K.L., K.O., and S.E.N. designed research; S.B., M.K.B., S.S., M.T., R.A., L.L., K.S., and S.E.N. performed research; S.B., M.K.B., R.A., R.O., M.L., K.S., and S.E.N. analyzed data; S.B., M.K.B., and S.E.N. wrote the paper with contributions form K.S., M.K.L and R.O.; and S.E.N. and M.K.B., designed illustrations for figures. All authors reviewed the results and approved the final version of the manuscript.

## Acknowledgements

We thank the Neal lab members, Darwin Gomez, Carla Calvo-Tusell, Fiona Kearns, Xandra Nuqui, and Lorenzo Casalino for in depth discussions and technical assistance, and Friederike Korn with the design of the TMED7-derived peptidyl α-ketoamides. These studies were supported by NIH grant 1R35GM133565-01, Pew Biomedical Award, NSF CAREER, and Freeman Hrabowski Scholar Award to S.E.N. R21CA256460 to A.J.O., Czech Science Foundation grant (project no. 20-25331S) and Ministry of Education, Youth and Sports (MEYS) of the Czech Republic InterCOST grant (project no. LUC23180) to K.S., and the project grant C10 from the Center for Molecular Medicine Cologne (CMMC) to M.K.L. Supercomputing resources were provided by XSEDE (NSF TG-CHE060073) and ACCESS (NSF TG- CHE060063) allocations and were granted to R.E.A.

## Declaration of Interests

The authors declare they have no competing interests within the contents of this article.

## Resource Availability

Further information and requests for resources and reagents should be directed to and will be fulfilled by the Lead Contact, Sonya Neal.

## Materials availability

Plasmids generated in this study is available from our laboratory.

## Data and Code Availability

Trajectories, structures, simulations scripts, analysis scripts, and data files are uploaded in Github page to enable other groups to use and explore the data we generated in this work. MD simulations dataset and analyses can be found in this Github link: https://github.com/mackevinbraza/RHBDL4_MD_simulations_dataset

## Experimental Procedures

### Cloning

The RHBDL4 gene was cloned into pcDNA3.1-HisA vector (Thermo Fisher Scientific) containing a hexahistidine-tag. The S144A mutant was created using site-directed mutagenesis (QuikChange II site-directed mutagenesis kit from Agilent) with the forward primer 5’- AAGGAGCTGTGCTGTAGGTTTCGCTGGAGTTTTGTTTGCTTTGAAA-3’ and reverse primer 5’-TTTCAAAGCAAACAAAACTCCAGCGAAACCTACAGCACAGCTCCTT-3’. The RHBDL4 mutant gene, W65A and G198A, were commercially synthesized by Genscript. Plasmid encoding human wild-type RHBDL4 with a C-terminal GFP had been described previously (Knopf et al. 2020). See Table S2 for all plasmids used in this study.

### *Mammalian Freestyle^TM^*Freestyle 293-F cell culture

pcDNA3.1-HisA plasmids expressing wild-type RHBDL4, RHBDL4-S144A, RHBDL4-W65A, and RHBDL4-G198A were transfected into Freestyle^TM^ 293-F mammalian suspension cells (Thermo Fisher Scientific) in Gibco Freestyle^TM^ 293-F media, according to the specified manufacturer’s protocol. pEGFP-N1-RHBDL4 was transfected as a 1:10 mix with empty pcDNA3.1 plasmid into Hek293T cells (ATTC) using 25-kDa linear polyethylenimine (Polysciences) as had been described (Knopf et al. 2020).

### Analysis of RHBDL4 expression levels

For analyzing RHBDL4 expression, 2×10^6^ A549 or 293-F cells in suspension were harvested and resuspended in lysis buffer (RIPA buffer, 0.01 M EDTA, and MgCl2). The samples were incubated for 30 min on ice, followed by addition of 1×Laemmli buffer at a 1:1 ratio. The sample was heated at 65°C for 10 min before being loading onto 12% SDS-PAGE gel. To check for endogenous RHBDL4 expression, western blotting was performed using a polyclonal α-Rhbdl4 (Sigma) or a monoclonal α-His antibody (Abclonal).

### RHBDL4 purification

Cells were harvested and resuspended in Buffer A (50 mM Hepes, pH 7.4, 500 mM NaCl, 10% (v/v) glycerol) and protease inhibitor cocktail (Sigma). For lysis, cell suspension was subjected to 15 strokes in an ice-cold, tight-fitting Dounce homogenizer (Pestle type B). The membrane fraction was pelleted by ultracentrifugation at 45,0000 rpm at 4°C for 30 min. The membrane pellet was resuspended in Buffer A supplemented with protease inhibitor cocktail and 3% (w/v) DDM and incubated at 4°C for 3 h. The insoluble material was ultracentrifuged at 95,000 rpm at 4°C for 3 h. HisPur Ni-NTA agarose resins (Thermo Fisher Scientific) was prepared by washing with water followed with Buffer B (50 mM Hepes, pH 7.4, 500 mM NaCl, 10% (v/v) glycerol, 0.1% (w/v) DDM) supplemented with 10 mM imidazole. Supernatant fraction containing solubilized membrane fraction was added to washed Ni-NTA resin and nutated at 4°C for 2 hr.

Protein-bound resin was transferred to a gravity flow column and washed with equal volume of Buffer B supplemented with 10 mM imidazole and Buffer B supplemented with 25 mM imidazole to wash away any unbound material. Additionally, beads were washed with 50 mM Hepes, pH 7.4, 450 mM NaCl, 50 mM KCl, 10% (v/v) glycerol, and 3 mM ATP to release any bound chaperones. Protein fractions were eluted with Buffer B supplemented with 500 mM imidazole. After removal of tag with TEV protease, the eluate was buffer-exchanged to 50 mM HEPES pH 7.4, 250mM NaCl, 10% (v/v) glycerol, and 0.1% (w/v) DDM, and concentrated using an Amicon filtration device with a 30-kDa molecular weight cutoff. Concentrated eluted proteins were further separated by size exclusion chromatography using a Superdex-75 10/300 GL column (Cytiva) equilibrated with 50 mM HEPES pH 7.4, 250mM NaCl, 10% glycerol, and 0.1% DDM. Protein concentration was determined by the BCA assay (Pierce BCA Protein Assay Kit, Thermo Fisher Scientific). The purified protein was aliquoted, flash frozen, and stored at - 80° C.

### FRET-based cleavage assays with IQ substrates

The enzymatic assay was performed as previously described (Arutyunova et al., 2014, Arutyunova et al., 2018). Briefly, 1.5 µM of wild-type RHBDL4 and inactive RHBDL4-S144A were incubated with 10 μM of IQ substrates at 37°C for 2 h in activity buffer containing 20 mM citrate phosphate pH 8.0, 150 mM NaCl, 20% (v/v) glycerol, and 0.1% (w/v) DDM. The maximum activity was obtained with the IQ6 peptide bearing the sequence Mca-Arg-Pro-Pro-Gly-Phe-Ser-Ala-Phe-Lys (DNP)-OH. The FRET-based cleavage assay was conducted in a fluorescence multi-well plate reader (SynergyMx, BioTek, VT, USA). Upon excitation at 320 nm, the emission intensity at 400 nm was recorded for 2 h at minimal intervals under shaking conditions and plotted using the GraphPad PRISM software. The initial velocities were defined as the slopes of the curve of the generated product as a function of time. The plots of the generated product against the corresponding substrate concentration were fitted with the Michaelis–Menten equation. Catalytic parameters were extrapolated using GraphPad PRISM. For determining cleavage sites, 1.5 µM of wild-type RHBDL4 and inactive RHBDL4-S144A were incubated with 10 μM of IQ6 or KSp207 internally quenched substrates for 4 h at room temperature, and the reaction was quenched by the addition of 8 M GuHCl. The samples were desalted using C18 tips (Rainin, Columbus, OH, USA), eluted in 80% acetonitrile, and dried down in a vacuum centrifuge prior to mass spectrometry analysis.

### Calculation of K_m_, V_max_, and K_cat_

All the calculations for K_m_, V_max_, and K_cat_ for RHBDL4 and its mutants were performed using GraphPad PRISM.

### Activity assay using a serine protease activity probe on recombinant RHBDL4

ActiveX™ TAMRA-FP serine hydrolase probe (Thermo Fisher Scientific) was resuspended in DMSO according to the manufacturer’s instructions. One hundred micrograms of wild-type RHBDL4 and inactive RHBDL4-S144A were incubated with 2 µM probe at room temperature for 1 h. The reactions were stopped by addition of 1×Laemmli buffer and heating the samples at 65°C for 10 min to deactivate active protease. The samples were subsequently run on a 12% SDS-PAGE gel and analyzed by western blotting using the Cy3 fluorescence filter of a Bio rad gel imaging system.

### Competition label assay in ER-enriched microsomes

ER-enriched microsomes were obtained by hypotonic swelling and centrifugation from Hek293T cells 24 h after transfection. To this end, cells from a sub-confluent 15 cm plate were resuspended in isolation buffer (10 mM HEPES-KOH pH 7.4, 1.5 mM MgCl2, 10 mM KCl, EDTA-free complete protease inhibitor cocktail from Roche). After 10 min incubation at 4°C, cells were lysed by passing six times through a 27-gauge needle. Cellular debris and nuclei were discarded after centrifugation at 1,000 g for 5 min at 4°C. The supernatant was spun at 100,000 g for 30 min at 4°C. The membrane pellet was resuspended in 40 µL RM buffer (50 mM HEPES-KOH pH 7.4, 250 mM sucrose, 50 mM KOAc, 2 mM Mg(OAc)2). For the competition label assay, 10 µL microsome suspension was either preincubated with the tested compound or the DMSO vehicle control for 1 h at 37°C. Subsequently, 500 nM TAMRA-FP serine hydrolase probe was added and incubate for 2 h at 37°C. The reaction was stopped by adding in 700 µl ice cold solubilization buffer (50 mM HEPES-KOH pH 7.4, 150 mM NaCl, 2 mM Mg(OAc)2, 10% (v/v) glycerol, 1 mM EGTA) containing 1% (w/v) Triton X-100. After incubation on ice for 30 min, RHBDL4-GFP was isolated by GFP-specific single-chain antibody fragment coupled to Sepharose beads (Fleig et al., 2012). After 1 h, immunoprecipitates were washed three times in solubilization buffer containing 0.5 % (w/v) Triton X-100 and then resuspended in SDS sample buffer followed by analysis on a 14 % SDS-PAGE gel. TAMRA-FP-labelling was analyzed in the wet gel with a green light source (535 nm) and a Cy3 filter by detecting fluorescence emission using ImageQuant 800 Fluor CCD imager (Cytiva) followed by western blotting using a α-GFP antibody (Roche).

### Peptide modeling and molecular dynamics simulations

To prepare the peptide model, we used AMBER tleap sequence module. The resulting model were solvated in the CHARMM-GUI server with Solution Builder.

### RHBDL4 modeling and molecular dynamics simulations

In the absence of resolved structure for human RHBDL4, we use the AlphaFold 2 model (UniProt ID Q8TEB9). We removed the low pLDDT scored-unstructured C-terminal domain residues (1 to 9) and N-terminal domain residues (221 to 315) to avoid highly fluctuating protein residues during the simulations. VBM and UIM binding motif was truncated due to their poorly scored structural model. Protonation state of the RHBDL4 residues at physiological pH 7.0 were assigned using PROPKA embedded in the CHARMM-GUI web server. All histidine residues are characterized with protonated N atom (i.e. HSD). Using CHARMM-GUI Membrane Builder, we embedded the protein in the ER membrane characterized with 15% cholesterol, 47% POPC [1- palmitoyl-2-oleoyl-glycero-3-phosphocholine], 20% POPE [1-palmitoyl-2-oleoyl-sn-glycero-3- phosphoethanolamine], 11% POPI [1-palmitoyl-2-oleoyl-sn-glycero-3-phosphoinositol], and 7% POPS [1-palmitoyl-2-oleoyl-sn-glycero-3-phospho-L-serine]. A water box with 30 Angstrom length size and a 0.15 M KCl were added (137 K^+^ and 67 Cl^-^ ions) to the membrane-protein-solvent system. CHARMM-GUI’s default NAMD configuration files for running membrane proteins with CHARMM force field was used in running molecular dynamics simulations. CHARMM36m force field was used for the protein and membrane while TIP3P force field was used for water. Temperature was set to 310 K and pressure at 1.01324 bar. The system was minimized for 5*x*10^6^ steps (with 1 fs/step and for a total of 5 ns) and then followed by several equilibration steps with varying constraint values. For the minimization, a 1 fs/step was used and then it was updated to 2 fs/step for the succeeding equilibration runs. The second equilibration lowers the constraintScaling from 10 to 5 and was equilibrated with 90x10^6^ steps. The third equilibration step used constant pressure where Barostat was set to 1.01325 bar using Langevin piston while the constraintScaling is set to 2.5. Fourth equilibration, constraintScaling was set to 1.0 and the timestep was increased to 2 fs/step. Fifth equilibration’s constraintScaling was set to 0.5 and sixth equilibration constraintScaling was set to 0.1. All the equilibration steps were done with anisotropic cell allowing the height, length, and width of the cell change independently. Particle Mesh Ewald (PME) was used to calculate long-range electrostatic interactions with PME grid spacing set to 1.0. Production was run with NPT ensemble by using 1.01325 bar pressure and 310 K temperature by using 2 fs/step timestep. Temperature was control with Langevin Dynamics. NAMD3 with CUDA was maximized by using CUDASOAintegrate with margin set to 4.

Velocities were set randomly from a Boltzmann distribution and differs in each replicate. Constant temperature was set to 310 K with Langevin dynamics. Pressure was control with Nose-Hoover Langevin piston. Furthermore, A 12 Angstrom cutoff was used to calculate electrostatic interactions with Particle Mesh Ewald. The scaled1-4 was used for the nonbonded parameters. NAMD3 MD engine was used in minimization, equilibration and production steps using multiple NVIDIA A40 GPUs. All the simulations were performed on the Delta supercomputer at the National Center for Supercomputing Applications (NCSA) - University of Illinois Urbana-Champaign. All-atom MD simulations of each system was run for 1000 ns (with n=4). This work gathered an accumulated 16,000 ns data of MD simulations of different protein-membrane-peptide systems.

Data analyses were done using cpptraj, pytraj, MDAnalysis, and matplotlib. All structures and MD simulations trajectories were rendered using VMD.

### Structure Clustering and Ensemble Docking

For both the RHBDL4 and peptide, we clustered the structures from the all-atom MD simulations using AMBER cpptraj. Trajectories were aligned from the first frame of the peptide. For the subsequent MD simulations of the RHBDL4-peptide systems, we used an open active site state of the RHBDL4 due to the fact that most of the clustered structures were closed. We defined the open active site as having at least one amino acid residue away from the RHBDL4 S144 and H195 (∼10 Angstrom). Top four clustered structures of the peptide PGFSA, QMESA, and MESA were docked using AutoDock Vina and AutoDock Tools. Docking was limited to the active site with grid box covering the crevice adjacent to the S144 and H195 residues.

### RHBDL4-peptide modeling and molecular dynamics simulations

To prepare the peptide model, we used AMBER tleap sequence module. The resulting model were solvated in the CHARMM-GUI server with Solution Builder. All the subsequent RHBDL4- peptide models were modeled and simulated similar to the RHBDL4 (apo) reported above.

---

**Supplemental Figure 1.**
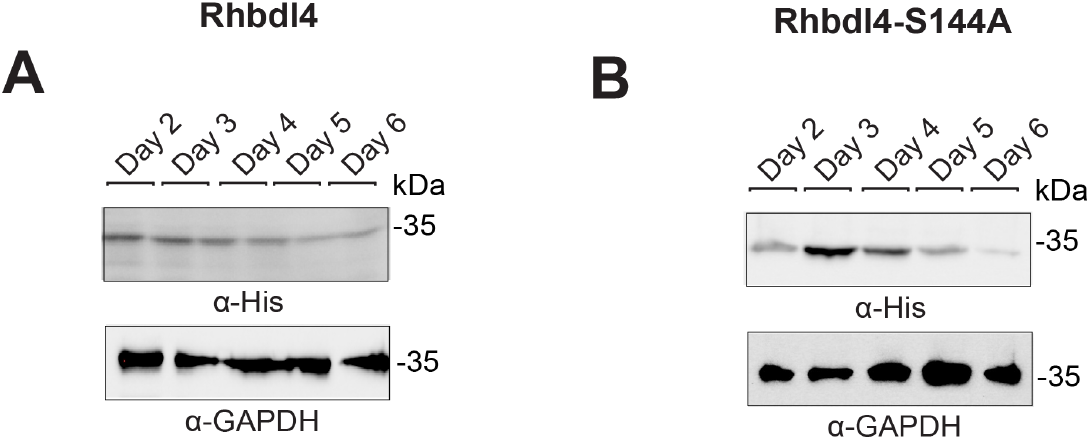
(A) Freestyle 293-F suspension cells were transfected with the RHBDL4 plasmid and cells were grown for the indicated number of days. 50 µg of lysate was subjected to immunoblotting for polyclonal α-RHBDL4 antibody and monoclonal α-GAPDH antibody. **(B)** Same as (A) except RHBDL4-S144A was transfected.

**Supplemental Figure 2.**
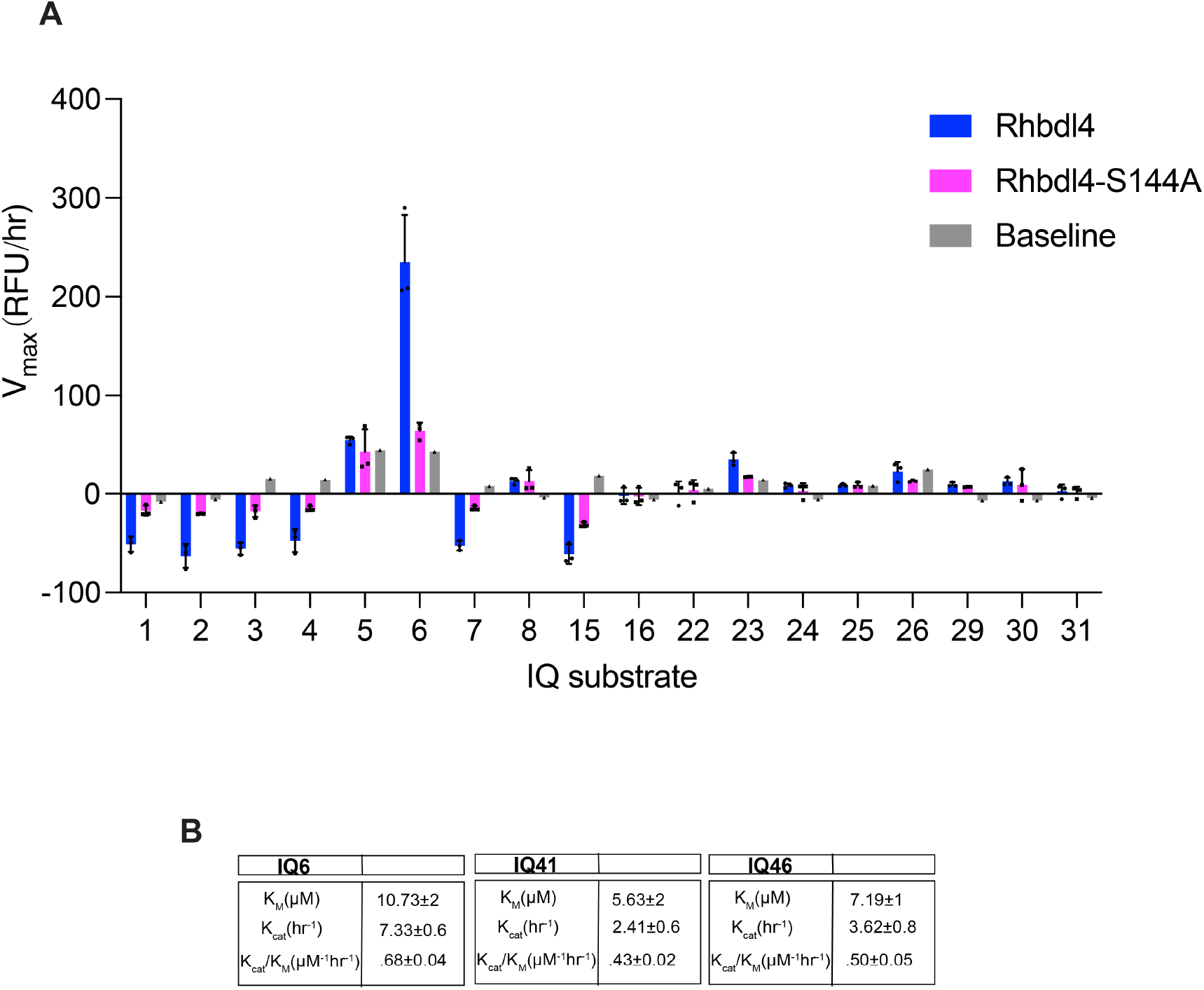
(A) Proteolytic rate of indicated IQ substrates cleaved by RHBDL4. **(B)** Catalytic parameters of IQ6, IQ40, and IQ41 substrate cleavage by RHBDL4.

**Supplemental Figure 3.**
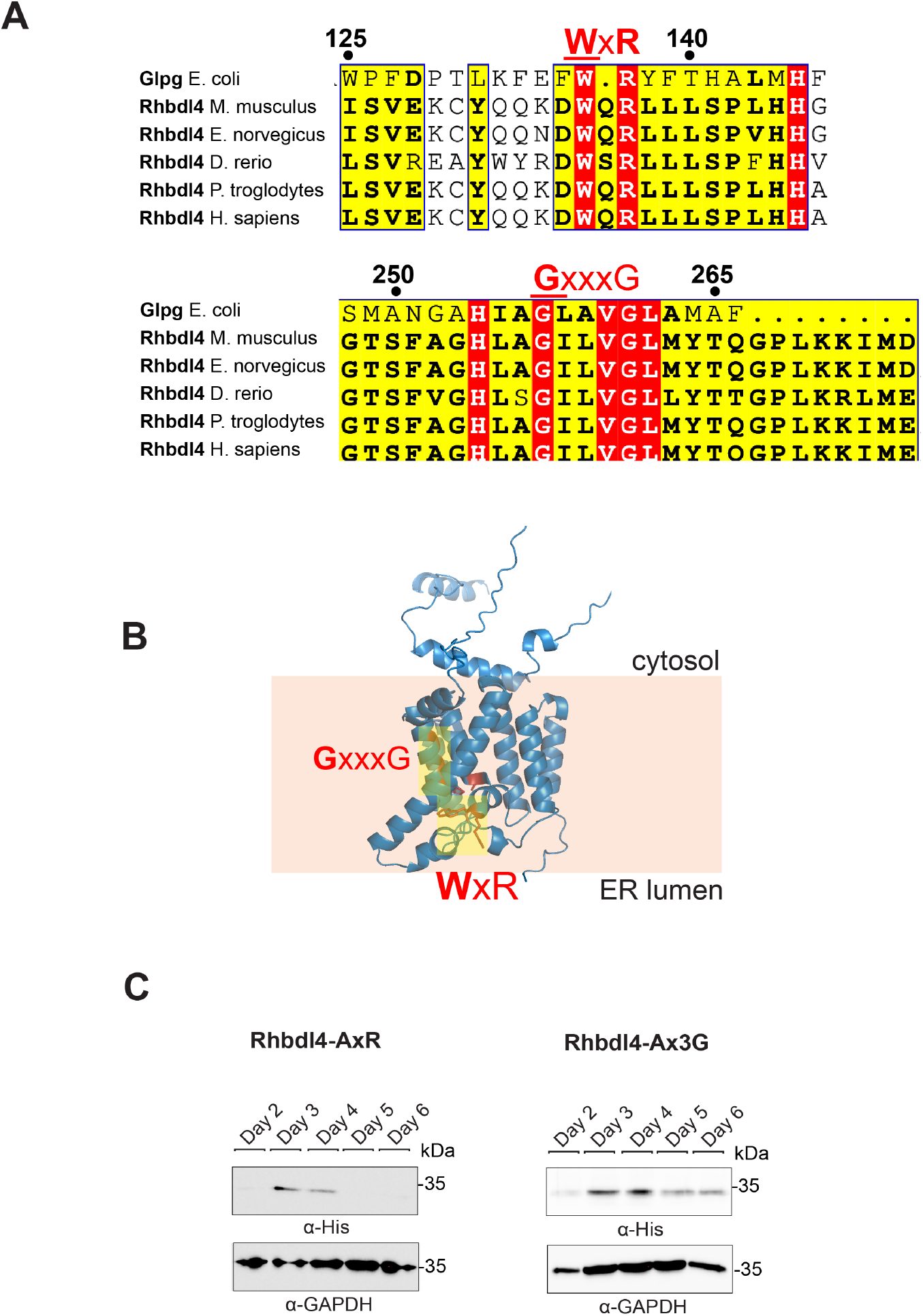
(A) Conservation of RHBDL4. Alignment of RHBDL4 (*H. sapiens, M. musculus, E. norvegicus, D. rerio, P. troglodytes*) with GlpG (*E. coli*). Identical residues are highlighted in red. **(B)** Alphafold model of RHBDL4. Positions of WxR and Gx3G motif is highlighted in yellow. **(C)** Freestyle 293-F suspension cells were transfected with RHBDL4-AxR and RHBDL4-Ax3G and cells were grown for the indicated amount of days. 50 µg of lysate was subjected to immunoblotting for RHBDL4 with monoclonal α-His antibody and monoclonal α- GAPDH antibody.

**Supplemental Figure 4.**
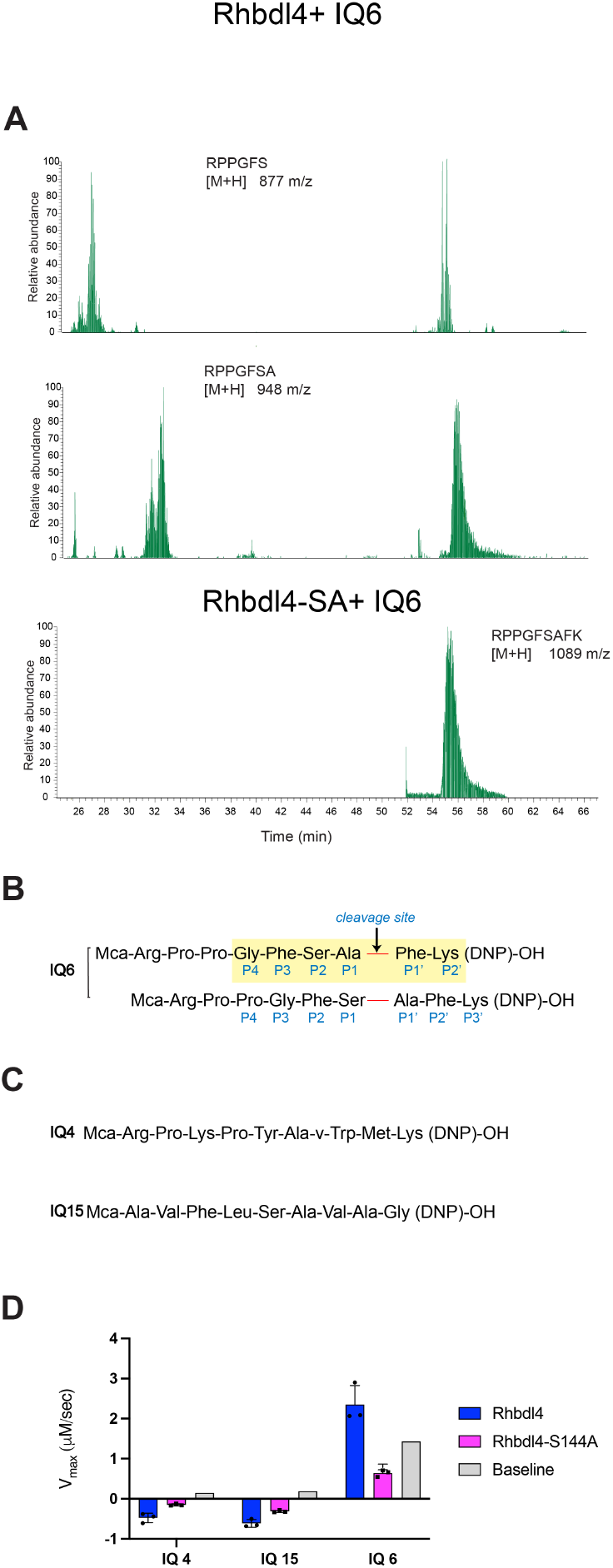
(A) Wild-type RHBDL4 and inactive RHBDL4-S144A were incubated with 10 μM of the IQ6 substrate Mca-Arg-Pro-Pro-Gly-Phe-Ser-Ala-Phe-Lys (DNP)-OH for 4 h at room temperature, and the reaction was quenched with 8 M GuHCl, desalted, eluted, and analyzed via tandem mass spectrometry (MS/MS). **(B)** Depiction of IQ6 cleavage sites as determined from tandem mass spectrometry (MS/MS). **(C)** Peptide sequence of GlpG-derived and PARL-derived fluorescent substrates IQ4 and IQ15, respectively. **(D)** Activity profiles of the wild- type RHBDL4 and mutant RHBDL4-S144A, as measured by cleavage of the fluorescent substrates IQ4 and IQ15. Assays was performed in triplicate, and data points are represented by the average ± standard deviation

**Supplemental Figure 5.**
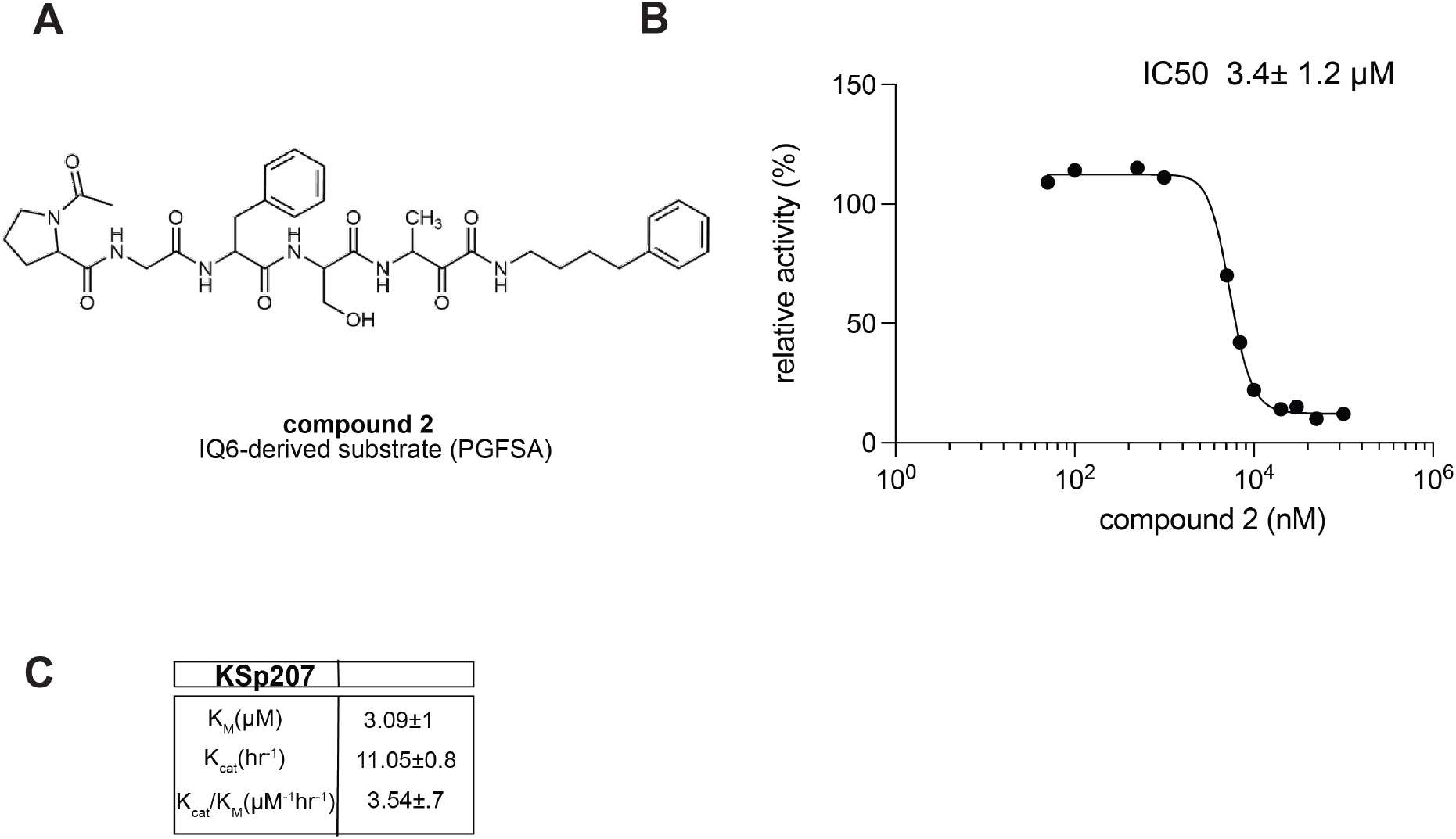
(A) Schematic representation of the peptidyl- α-ketoamide inhibitor, compound **2. (B)** Representative inhibition curve derived from measuring rates of RHBDL4 proteolysis of the fluorescent substrate IQ6 with increasing concentrations of compound **2**. **(C)** Catalytic parameters of KSp207 substrate cleavage by RHBDL4.

**Supplemental Figure 6.**
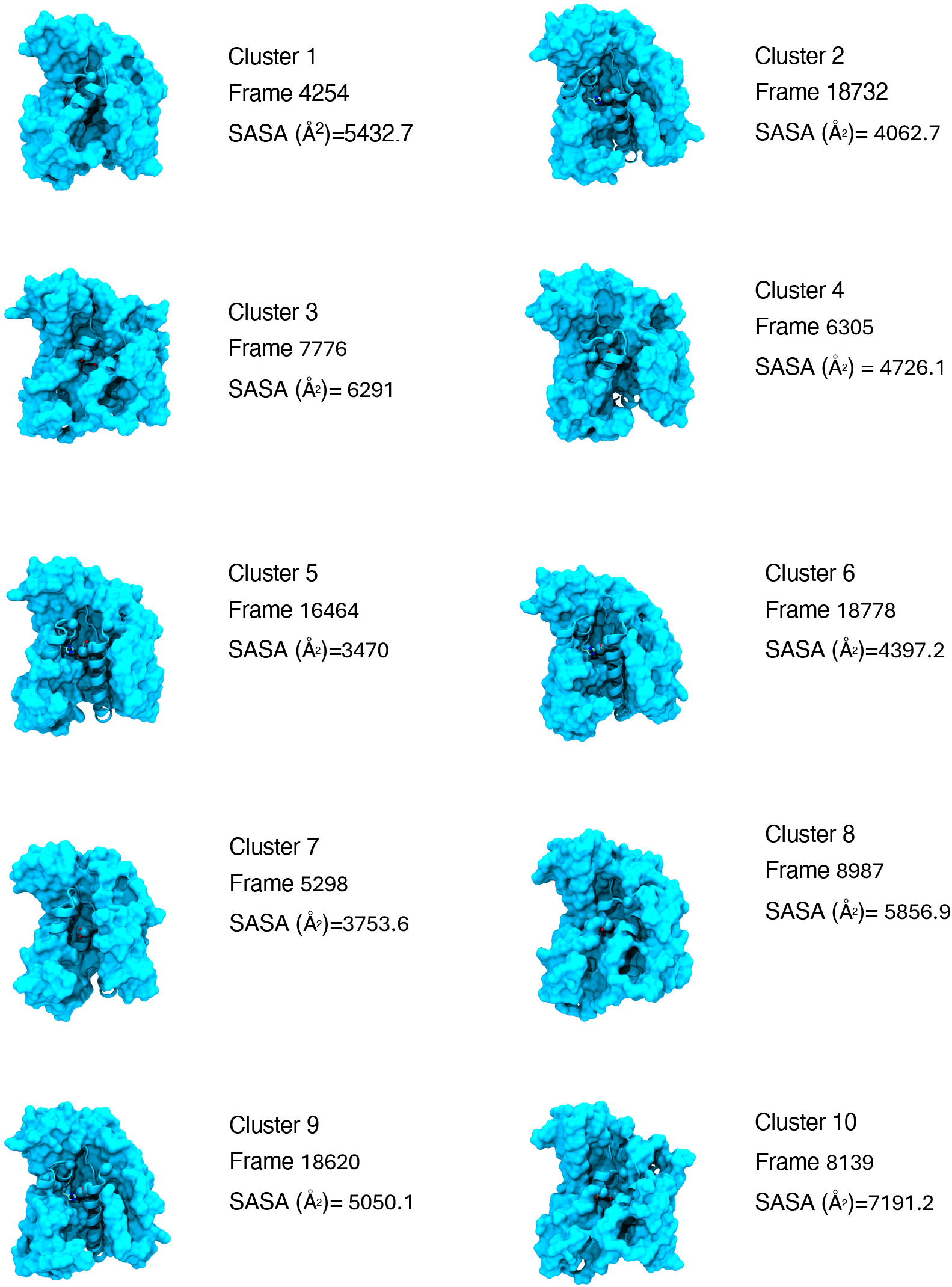
RHBDL4 (apo) most populated clusters from all-atom MD simulations. Structures were clustered based from the solvent-accessibility-surface-areas (SASA) values of residues within 10 Angstrom of HIS144 and SER195. Each cluster was represented with the frames that has max SASA values.

**Supplemental Figure 7.**
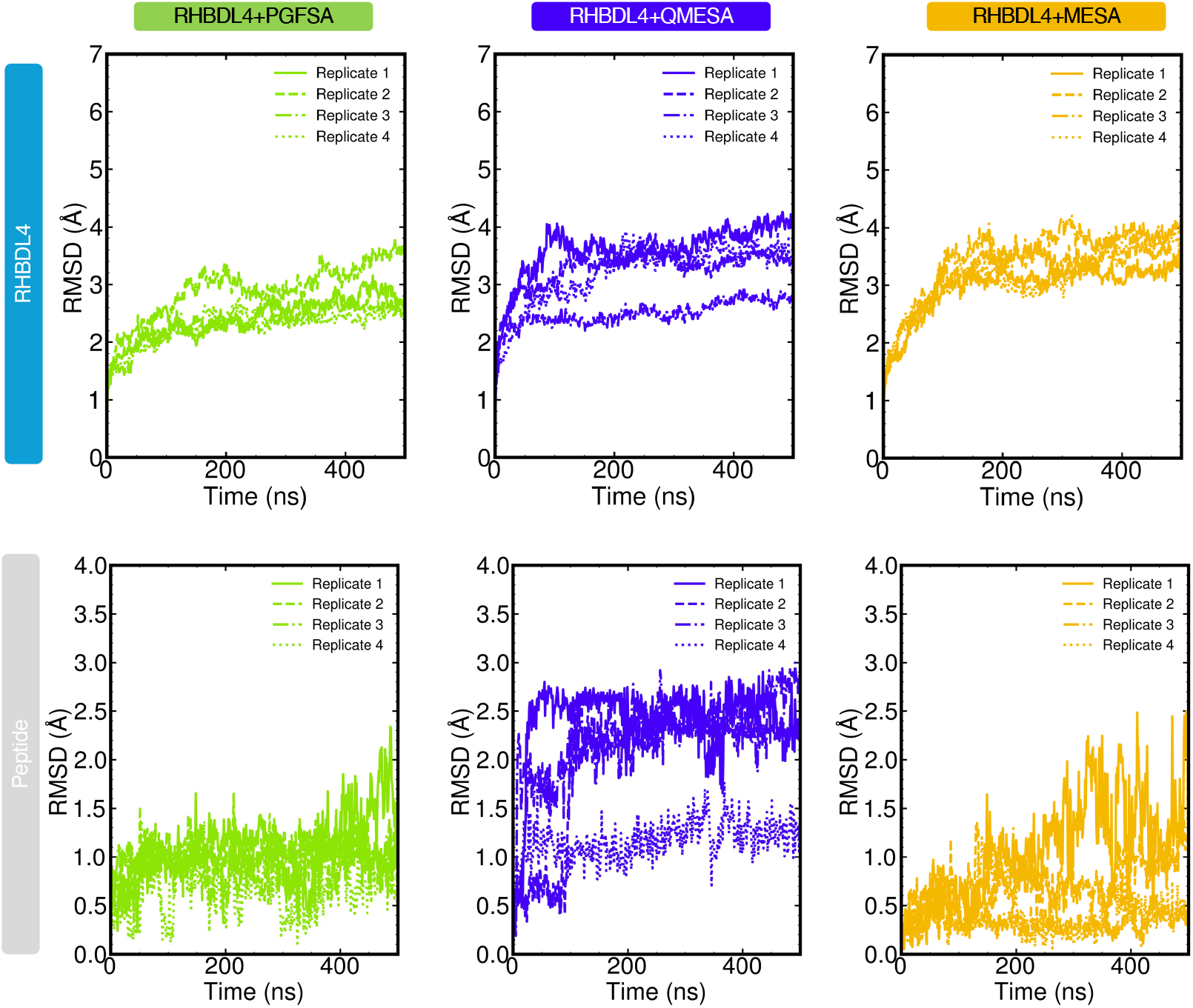
RHBDL4 and peptide Ca Root-mean-square deviation from the initial frame of molecular dynamics simulations. Trajectories were aligned. Four replicates for each system were analyzed.

**Supplemental Figure 8.**
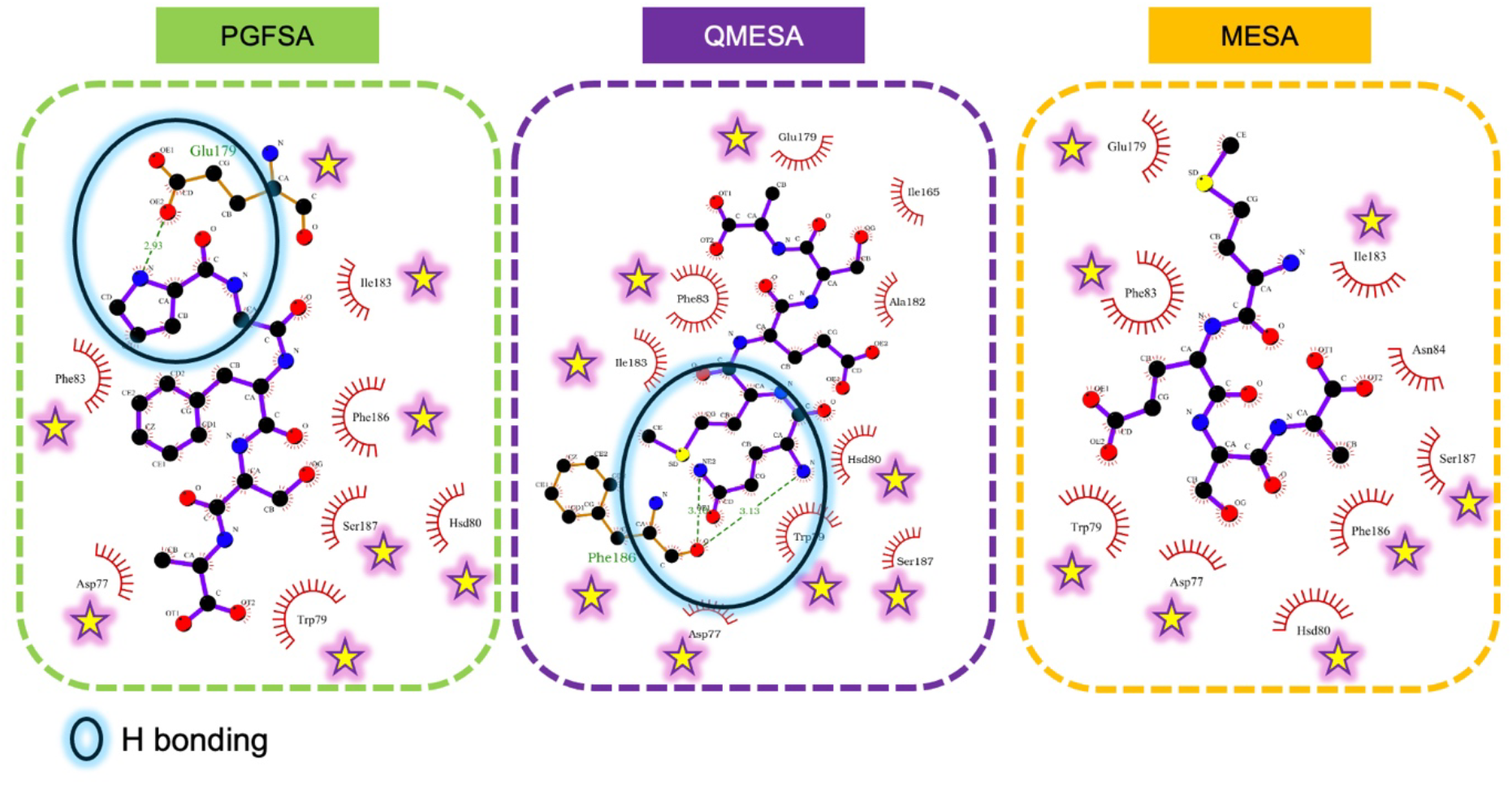
2D representation of RHBDL4 close interacting residues with the peptide side chains. All residues that were shared by the three substrate-derived peptides were marked with yellow stars. Ligplot+ was used to render this figure.

**Supplemental Figure 9.**
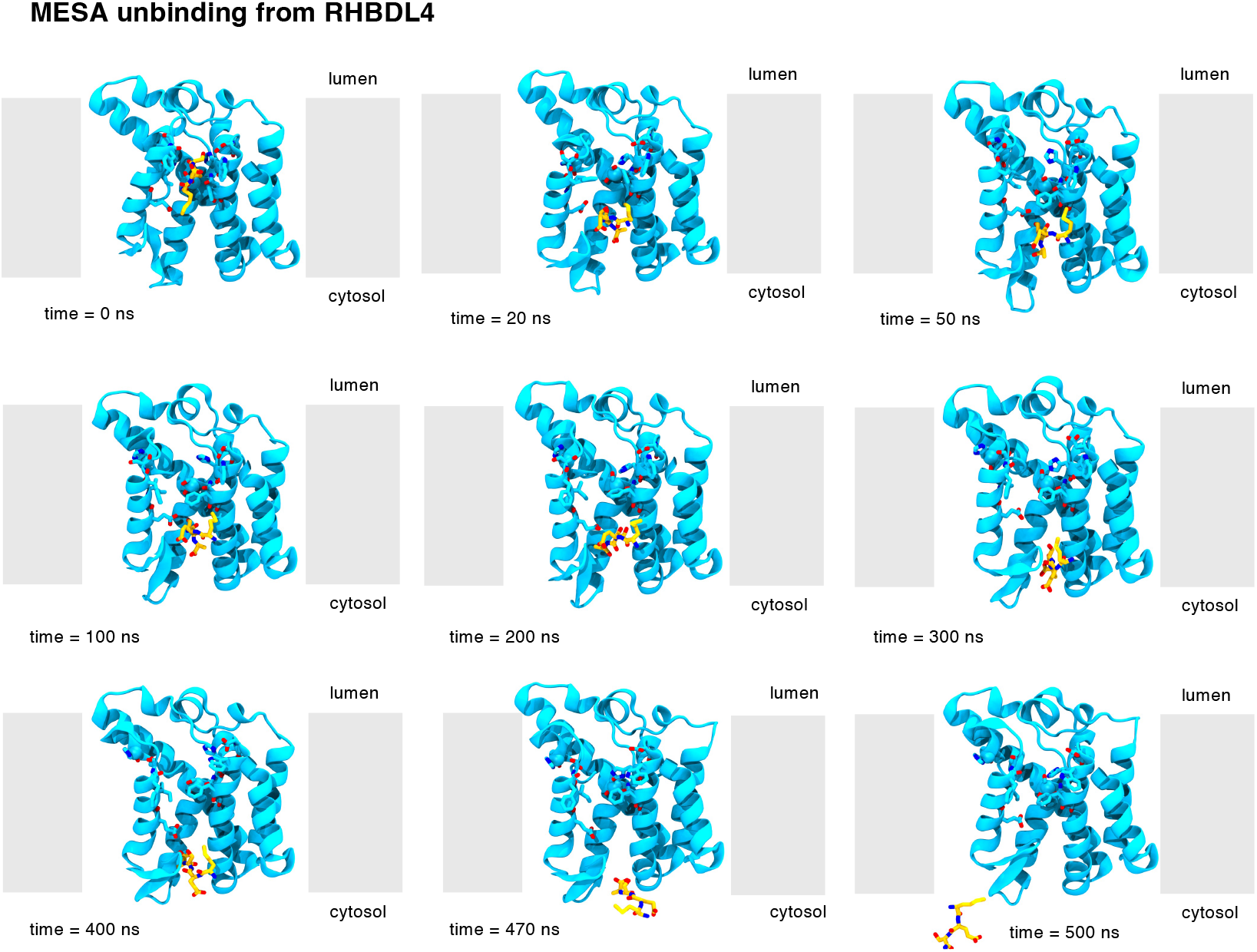
Time evolution snapshot of MESA interaction with the RHBDL4 active site.

**Supplemental Figure 10.**
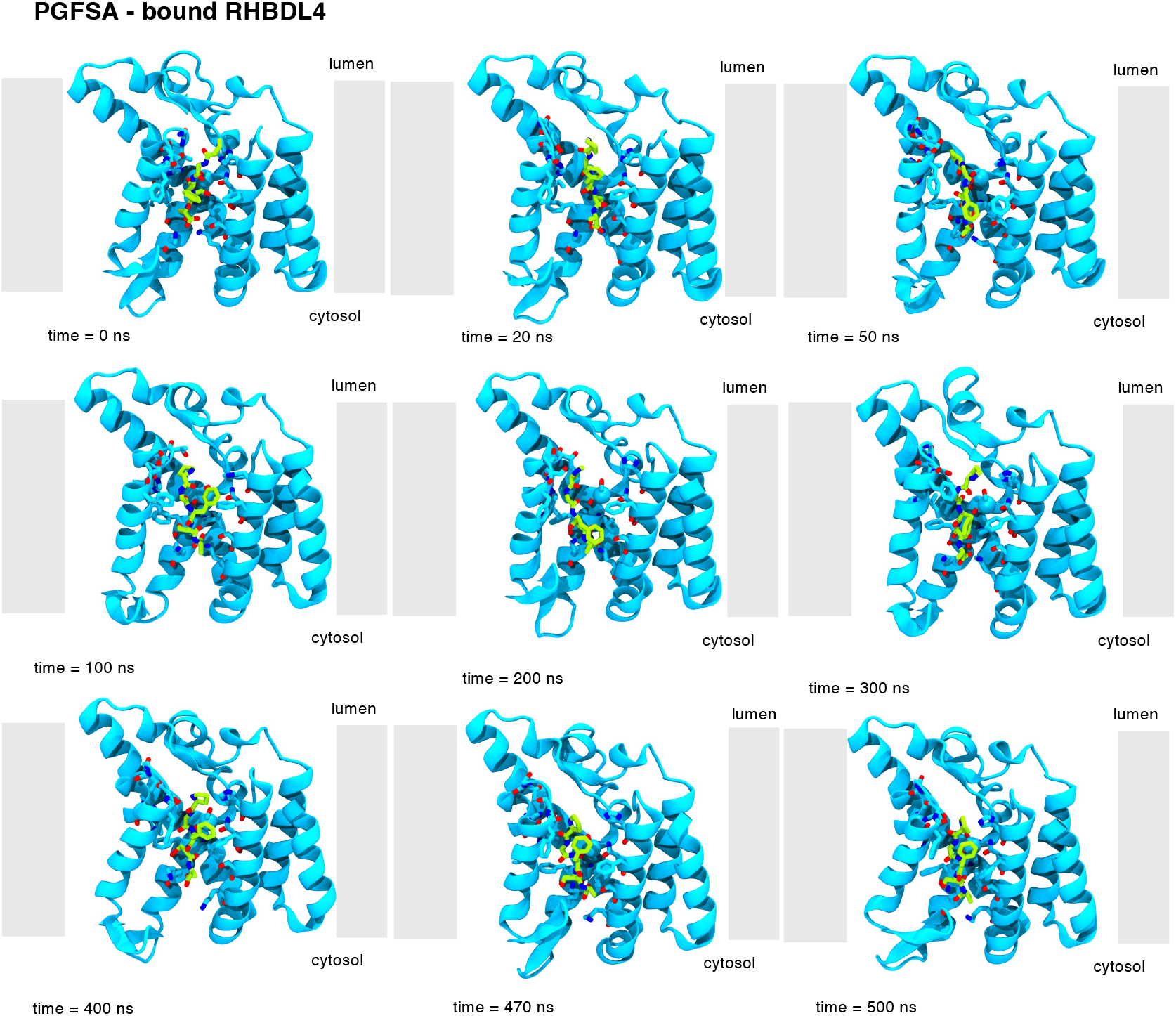
Time evolution snapshot of PGFSA interaction with the RHBDL4 active site.

**Supplemental Figure 11.**
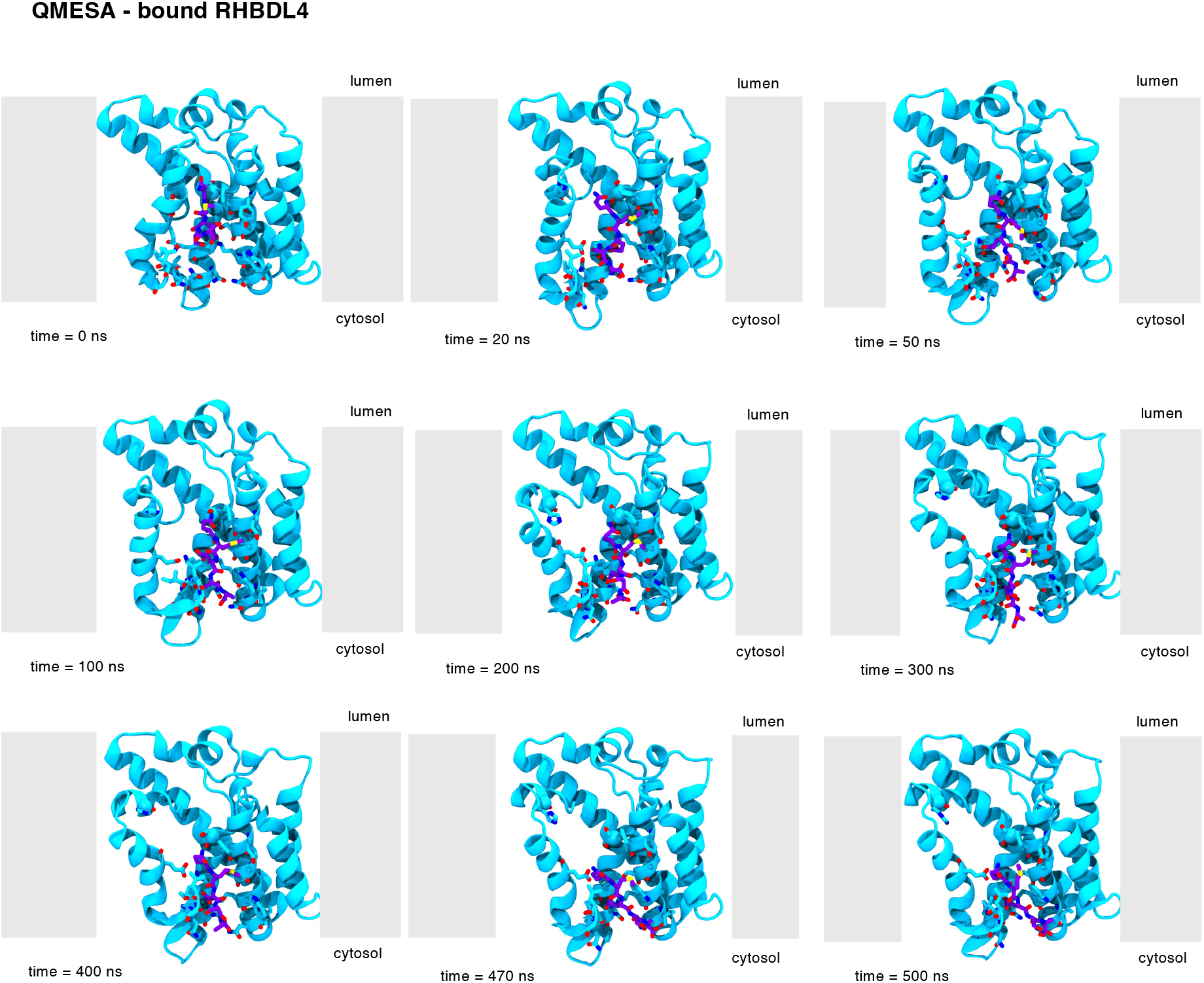
Time evolution snapshot of QMESA interaction with the RHBDL4 active site.

**Supplemental Table 1:** AutoDock Vina ensemble molecular docking (n=4)

|  | **PGFSA (IQ6)**<br>mca-<br>RPPGFSAFK(dnp) | **QMESA (Ksp207)**<br>Ac-QMESA-4mc | **MESA** |
| --- | --- | --- | --- |
| **Binding affinity** (kcal×mol<sup>-1</sup>)* | -6.8 ± 0.5 | -5.2 ± 0.3 | -5.2 ± 0.1 |

**Supplemental Table 2.** Plasmids used in this study.

| **Plasmid #** | **Backbone & Gene** |
| --- | --- |
| pSN182 | pcDNA3.1 HisA RHBDL4 |
| pSN183 | pcDNA3.1 HisA RHBDL4-S144A |
| pSN294 | pcDNA3.1 HisA RHBDL4-AxR |
| pSN295 | pcDNA3.1 HisA RHBDL4-Ax3R |

## Notes

https://github.com/mackevinbraza/RHBDL4_MD_simulations_dataset

### Competing Interest Statement

The authors have declared no competing interest.

### Summary of Updates

Fixed and updated figures. Added a technical limitations section.

